# Caspase-8 Modulates Angiogenesis By Regulating A Cell Death Independent Pathway In Endothelial Cells

**DOI:** 10.1101/708651

**Authors:** Nathalie Tisch, Aida Freire-Valls, Rosario Yerbes, Isidora Paredes, Silvia La Porta, Xiaohong Wang, Rosa Martín-Pérez, Laura Castro, Wendy Wei-Lynn Wong, Leigh Coultas, Boris Strilic, Hermann-Josef Gröne, Thomas Hielscher, Carolin Mogler, Ralf Adams, Peter Heiduschka, Lena Claesson-Welsh, Massimiliano Mazzone, Abelardo López-Rivas, Thomas Schmidt, Hellmut G. Augustin, Carmen Ruiz de Almodovar

## Abstract

During developmental angiogenesis blood vessels grow and remodel to ultimately build a hierarchical vascular network. Whether and how cell death signaling molecules contribute to blood vessel formation is still not well understood. Caspase-8 (Casp-8), a key protease in the extrinsic cell death-signaling pathway, regulates both cell death via apoptosis and necroptosis. Here we show that expression of Casp-8 in endothelial cells (ECs) is required for proper postnatal angiogenesis. EC specific Casp-8 knockout pups (Casp-8^ECko^) have reduced retinal angiogenesis, as the loss of Casp-8 reduced EC proliferation, sprouting and migration independent of its cell death function. Instead, the loss of Casp-8 caused hyperactivation of p38 mitogen-activated protein kinase (MAPK) downstream of receptor-interacting serine/threonine-protein kinase 3 (RIPK3) and destabilization of VE-cadherin at EC junctions. In a mouse model of oxygen-induced retinopathy (OIR), resembling retinopathy of prematurity (ROP), loss of Casp-8 in ECs is beneficial, as pathological neovascularization was reduced in Casp-8^ECko^ pups. Taken together, we identify that Casp-8 signals in a cell-death independent manner in ECs during postnatal and pathological blood vessel formation.

## INTRODUCTION

Functional blood vessels are of vital importance. Impaired vessel formation contributes to many pathological situations, among others to ischemic or inflammatory disorders. Ischemic retinopathies are the main causes of severe visual impairment and sight loss in premature children, diabetic adults and the elderly population. For example, retinopathy of prematurity (ROP), an ocular neovascular disease and the major cause of acquired blindness in preterm infants (1), is characterized by excessive angiogenesis, breakdown of the endothelial barrier, vascular leakage, edema, hemorrhages, retinal detachment and compromised vision (1). Although certain strategies to block excessive angiogenesis exists (2), identification of other pathways that regulate physiological and pathological vessel growth is of high interest in order to further develop better or complementary therapeutic treatments.

The formation of functional blood vessel networks starts with the initial formation of a primitive vascular plexus from which new blood vessels sprout, coordinately expand and branch. Redundant vessel branches are then selectively removed by vessel pruning to ultimately establish a hierarchical vascular network (3, 4). Whereas apoptosis has been implicated in certain conditions of vessel remodeling (5–9) and in the regulation of capillary vessel diameter (10), it is not known whether other forms of cell death contribute to developmental angiogenesis or whether cell death signaling molecules have cell death-independent functions in endothelial cells (ECs).

Caspase-8 (Casp-8) is the initiator caspase of the extrinsic apoptosis pathway that activates the effector Caspase-3 (11). Cellular FLICE inhibitory protein (c-FLIP), an inactive homolog of Casp-8 that lacks its catalytic activity, can heterodimerize with Casp-8, prevent its full activation, and thus inhibit apoptosis (11). The c-FLIP/Casp-8 heterodimer still possesses a basal Casp-8 activity necessary to inhibit a second type of programmed cell death, called necroptosis (a non-apoptotic cell death characterized by swelling and rupture of the cell membrane) (12, 13). To do so, Casp-8 cleaves the receptor-interacting serine/threonine-protein kinase 3 (RIPK3) and therefore blocks the activation of the ultimate effector of necroptosis, the pseudokinase mixed-lineage kinase domain-like (MLKL), downstream of RIPK3 (13). Therefore, Casp-8 is a central signaling node in the decision of whether a particular cell dies by apoptosis or necroptosis. In addition to its role in cell death signaling, Casp-8 has been described to regulate cell death-independent processes such as cell migration (14, 15), or proliferation (16) and to act as DNA-damage sensor induced by cell proliferation (17).

Deletion of Casp-8 in mice is embryonically lethal. Interestingly, Casp-8 knockout mouse embryos die at mid-gestation presenting a circulatory failure phenotype and damaged blood vessel capillaries (18–20). Endothelial cell (EC) specific knockout of Casp-8 during embryonic development mimics the severity of the full knockout (21), suggesting that Casp-8 in ECs is required for the formation of a proper vascular system. However, whether Casp-8 is also required at later timepoints during vessel development, and whether it plays a cell-death dependent or independent function, remains unknown. In this regard, cell death-dependent and -independent functions of RIPK3 have been described in pathological angiogenesis. In a mouse model of melanoma metastasis, it has been shown that RIPK3 is involved in tumor-induced EC necroptosis but also in vessel permeability via activation of p38 (22, 23). Whether RIPK3 is not only contributing to pathological vessel formation and function, but also to vessel stability in physiological conditions of angiogenesis, is so far unknown.

Here, using a conditional inducible EC specific Casp-8 knockout mouse line, we show that the loss of Casp-8 results in reduced postnatal angiogenesis. Most strikingly, we demonstrate that this phenotype is independent of necroptosis, as the loss of Casp-8 on a MLKL-null background still results in vascular defects. Instead, Casp-8 regulates EC proliferation, sprouting and migration, as well as the stability of adherens and tight junctions. Mechanistically, we show that loss of Casp-8 leads to the constitutive phosphorylation of p38 mitogen-activated protein kinase (MAPK) via RIPK3 and to defects in VE-cadherin subcellular localization. Applying the oxygen induced retinopathy model (OIR), a model that resembles the pathology of ROP, we show that the loss of Casp-8 in ECs is beneficial, as pathological neovascularization in Casp-8 knockout mice is reduced.

## RESULTS

### Postnatal deletion of Casp-8 in ECs results in impaired angiogenesis in the retina

To explore the function of Casp-8 during developmental angiogenesis *in vivo*, we generated inducible EC specific Casp-8 knockout mice. To do so, we crossed the tamoxifen inducible Cdh5-(PAC)-CreERT2 mouse line (24), where Cre-recombinase is expressed under control of the EC specific Cdh-5 promoter, with Casp-8^fl/fl^ mice. Consistent with previously published data (21), therefore validating our generated mouse line, knockout of Casp-8 in the endothelium during embryonic development resulted in yolk sac and embryo vascular defects as well as in increased embryonic lethality and defects in the yolk sac vasculature (Suppl. Fig. 1A,B).

To analyze the effect of loss of Casp-8 in ECs during postnatal vascular development, we induced Cre recombination in newborn pups as previously described (25) (Fig. 1A) and checked the recombination efficiency in isolated lung ECs (Suppl. Fig. 1C,D). Tamoxifen treatment resulted in 70% reduction of Casp-8 mRNA levels in lung ECs from Casp-8^ECko^ pups at postnatal (P) day 6 (Suppl. Fig. 1D). In contrast to the lethal effects of loss of Casp-8 in the embryonic vasculature, postnatal knockout of Casp-8 was not lethal during the time the mice were observed (until P15) and pups were indistinguishable from Casp-8^WT^ littermates (Suppl. Fig. 1E-G). As Casp-8 is expressed in ECs of developing blood vessels in the retina (Suppl. Fig. 1H), and as this is a well-established model to postnatally study developmental angiogenesis (26), we used the growing retina for our purposes. Analysis of the vasculature (by staining with the endothelial marker IsoB4) in P6 retinas showed that the total area covered by blood vessels and the vascular outgrowth were reduced in Casp-8^ECko^ retinas compared to Casp-8^WT^ littermates (Fig. 1B-D). The number of vessel branches was also decreased (Fig. 1E,F), indicating a reduced complexity of the vascular network in Casp-8^ECko^ pups. In addition, we counted less sprouts at the angiogenic front in Casp-8^ECko^ pups (Fig.1 G,H).

**Figure 1.**
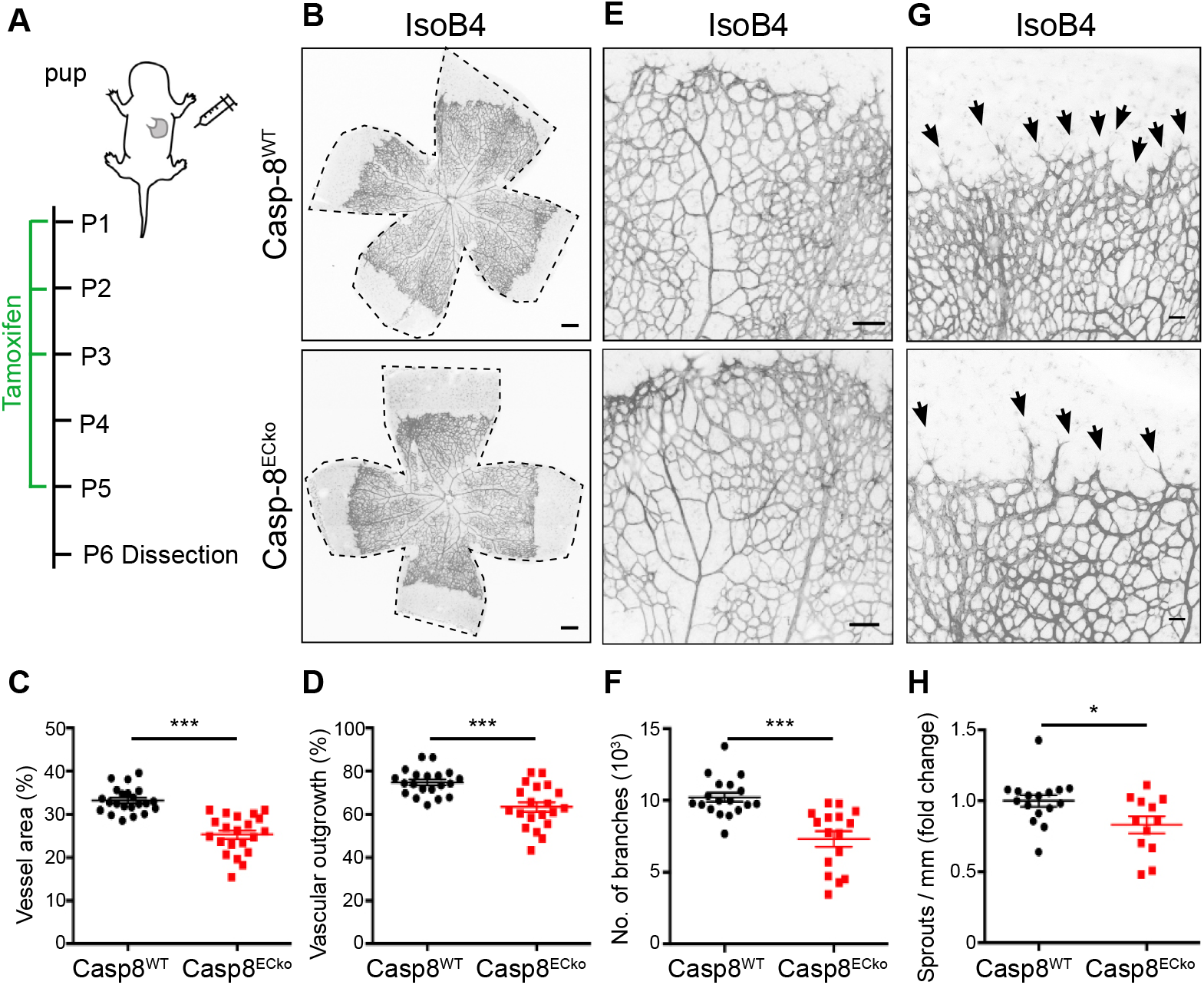
Postnatal EC specific knockout of Casp-8 results in impaired angiogenesis. **A)** Scheme of tamoxifen administration in pups. **B)** Representative images of whole mount P6 retinas stained with IsoB4 (ECs) in Casp-8^WT^ and Casp-8^ECko^ mice. Dashed black lines highlight the total retina area. **C,D)** Quantification of vessel area (C; n=22 WT, 21 ECko) and retina vessel outgrowth (D; n=20 WT, 20 ECko). **E)** Representative higher magnifications of the retina stained with IsoB4. **F)** Quantification of number of branches (n=18 WT, 16 ECko). **G)** Representative images of the retina angiogenic front stained with IsoB4. Black arrows point to EC sprouts. **H)** Quantification of the number of sprouts per front length showing reduced number of sprouts in Casp-8^ECko^ retinas (n=16 WT, 12 ECko). For C,D,F,H data represent mean ± SEM from 4 independent litters (* *P* <0.05, *** *P* <0.001; two-tailed unpaired Student *t*-test). Scale bars: B) 100μm; E) 50μm; G) 20μm.

Altogether, this data show that loss of Casp-8 in the endothelium results in vascular defects during developmental angiogenesis.

### Necroptosis does not contribute to the vascular defects in Casp-8^ECko^ pups

During postnatal angiogenesis in the retina, the vascular network develops to its final mature stage via a combination of different cellular processes such as EC proliferation, migration, vessel maturation and vessel remodeling. As Casp-8 regulates cell death and survival, and as this contributes to vessel remodeling and regression (5, 8, 9), we checked whether knockout of Casp-8 would result in EC death due to activation of the necroptotic pathway. *In vivo*, an indirect way to assess cell death is the quantification of regressing vessel branches, which leave a Collagen type IV (ColIV^+^) empty sleeves behind (27). Therefore, we analyzed vessel regression by quantifying the number of CollV^+^ IsoB4^−^ sleeves and found no differences between genotypes (Fig. 2A,B). In line with these results, pericyte coverage analyzed by co-staining of the pericyte marker Desmin and IsoB4 revealed no differences between Casp-8^ECko^ and Casp-8^WT^ retinas, indicating that vessel maturation and stabilization were also not affected (Fig. 2C, D). Interestingly, we noticed a small but significant decrease in the number of cleaved Caspase-3^+^ (cCasp3^+^) and TUNEL^+^ ECs in Casp-8^ECko^ pups (Fig. 2E-H), suggesting that Casp-8 mediated apoptosis via the extrinsic cell death signaling pathway may have a small but significant contribution to the physiological remodeling process, however without affecting overall vessel regression. To further explore this hypothesis, we first analyzed the ability of ECs to die by necroptosis *in vitro*. For this purpose, we counted the number of propidium iodide positive cells (PI^+^, which detects apoptotic and necroptotic cells (28)) in wildtype (Casp-8^WT^) and Casp-8 knockdown (Casp-8^KD^) HUVECs using a lentivirus–mediated shRNA knockdown system (29) (Suppl. Fig. 2A). Both Casp-8^WT^ and Casp-8^KD^ HUVECs showed a slight, though not significant, increase in the number of PI^+^ cells upon TRAIL or TNF stimulation (Suppl. Fig. 2B). As a positive control to demonstrate that HUVECs were not just resistant to cell death, we knocked down c-FLIP (c-FLIP^KD^), the intrinsic inhibitor of Casp-8. c-FLIP^KD^ ECs presented a significant increase in cell death upon TNF or TRAIL stimulation (Suppl. Fig. 2B). As vessel development in the retina is regulated by hypoxia (30), and hypoxia can be a modulator of cell death (31), we also performed the same experiments under hypoxic conditions, which produced the same results (Suppl. Fig. 2C).

**Figure 2.**
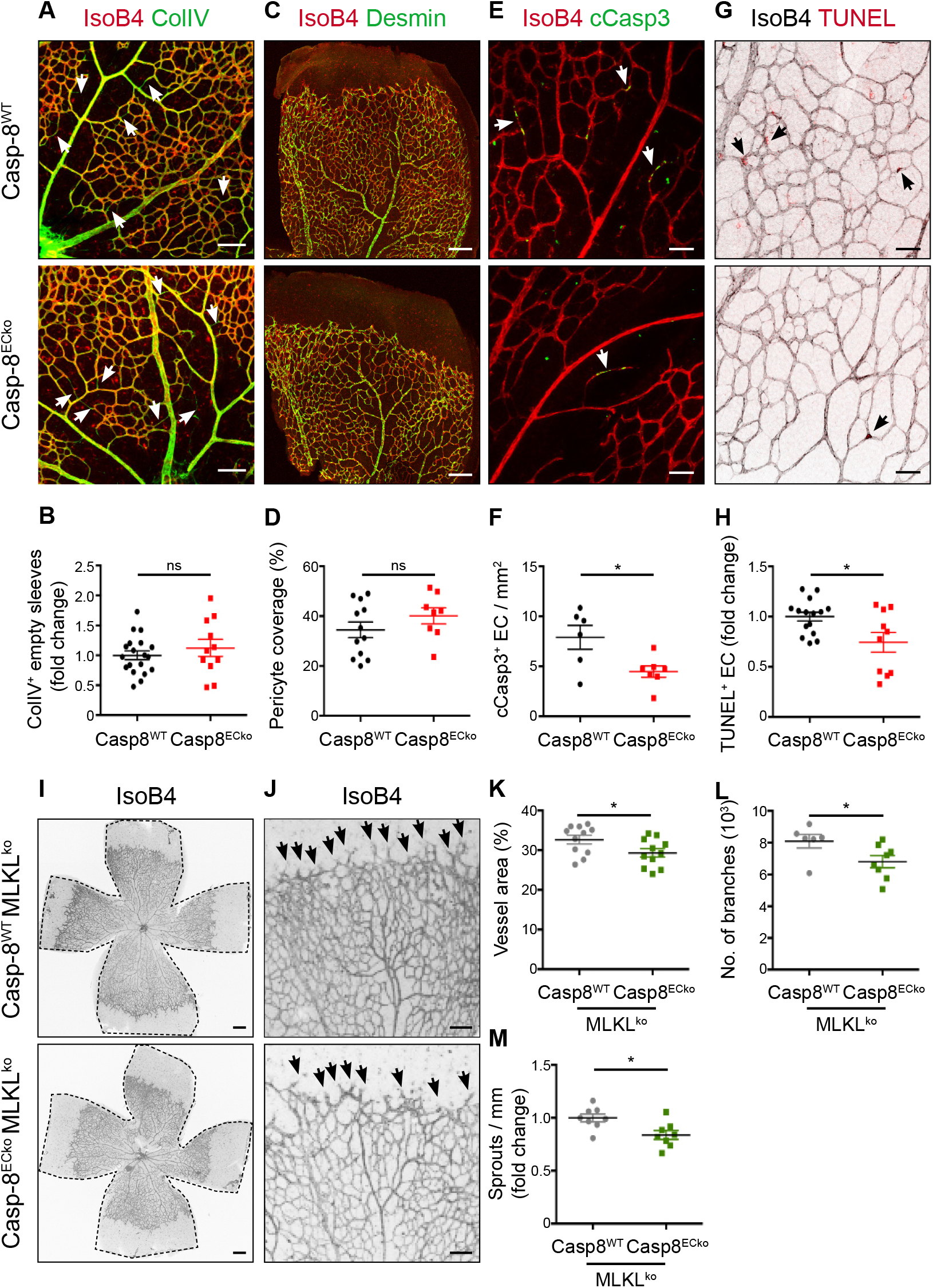
Loss of Casp-8 in ECs does not result in vessel regression nor necroptosis. **A)** Retinas from P6 pups were co-stained with IsoB4 and Collagen IV (ColIV) in Casp-8^WT^ and Casp-8^ECko^ retinas. **B)** Quantification of relative vessel regression shows no significant differences between genotypes (n=19 WT, 11 ECko). **C)** Representative images of pericyte coverage. Retinas were co-stained with Desmin (pericyte marker) and IsoB4. **D)** Quantification of Desmin^+^ area per vascular area (%) shows no significant differences between genotypes (n=12 WT, 8 ECko). **E)** Representative images of apoptotic ECs (white arrows), co-stained with IsoB4 and cleaved caspase-3 (cCasp3). **F)** Quantification of cCasp-3^+^/IsoB4^+^ cells per vessel area revealing fewer apoptotic ECs in Casp-8^ECko^ retinas compared to Casp-8^WT^ (n=6 WT, 7 ECko). **G)** Representative images of retinas co-stained with IsoB4 and TUNEL (black arrows point to TUNEL+ ECs). Images of IsoB4 were transformed to grey colors with ImageJ for better visualization. **H)** Quantification of relative amount of TUNEL^+^ ECs per vessel area also shows decreased numbers of apoptotic ECs in Casp-8^ECko^ retinas (n=15 WT, 10 ECko). **I,J)** Representative images of the retinal vasculature stained with IsoB4 in Casp-8^WT^/MLKL^ko^ and Casp-8^ECko^/MLKL^ko^ mice. Black arrows point to EC sprouts. **K-M)** Quantitative analysis showing reduced vessel area (K; n=11 WT, 11 ECko), number of branches (L; n=6 WT, 8 ECko) and reduced number of sprouts per front area (M; n=6 WT, 8 ECko) in Casp-8^ECko^/MLKL^ko^ retinas compared to Casp-8^WT^/MLKL^ko^ littermates. For B,D,F,H,K-M data represent mean ± SEM at least from 3 independent litters (* *P* <0.05, ns: not significant; two-tailed unpaired Student *t*-test). Scale bars: A,C,I) 100μm; E,G) 20μm; J) 50μm.

Finally, to determine whether necroptosis could contribute to the observed phenotype *in vivo*, we genetically deleted MLKL (the ultimate effector of necroptosis (13)) in the Casp-8^ECko^ mice by crossing Cdh5-(PAC)-CreERT2 x Casp-8^fl/fl^ mice with a MLKL^ko^ mouse line. The vessel area and the number of vessel branches at P6 were not affected in heterozygous or homozygous MLKL^ko^ pups compared to wildtype littermates (Suppl. Fig. 3A-D), indicating that MLKL alone did not contribute to angiogenesis. However, the vessel area and the number of vessel branches as well as the sprouts at the angiogenic front were still reduced in Casp-8^ECko^/MLKL^ko^ pups (compared to Casp-8^WT^/MLKL^ko^ pups) (Fig. 2I-M), showing that blocking necroptosis in Casp-8^ECko^ pups did not rescue the vascular defects.

Our data indicate that the loss of Casp-8 in ECs during postnatal development does not induce cell death via necroptosis. Even though we found a mild decrease in the number of cCasp-3^+^ and TUNEL^+^ ECs at P6 in Casp-8^ECko^ pups, this did overall not affect the percentage of vessel regression, indicating that extrinsic cell death is not an active driver of vessel pruning during postnatal vascular development. Our data further suggest that Casp-8 regulates developmental angiogenesis in a cell death-independent way.

### Casp-8 regulates VEGF-induced EC sprouting, proliferation, and migration

It has been shown that Casp-8 has diverse cell death-independent functions, for example in the regulation of cell migration (14, 32). Thus, to further characterize the EC phenotype and to understand which processes are impaired upon loss of Casp-8, we analyzed the response of Casp-8^KD^ ECs upon VEGF stimulation. To do so, we analyzed EC sprouting *in vitro* using the bead sprouting assay (33). While knockdown of Casp-8 (using a specific siRNA, Suppl. Fig. 4A) had no effect on unstimulated conditions, Casp-8^KD^ ECs failed to respond to VEGF stimulation (Fig. 3A,B). Additionally, we tested the response of Casp-8^KD^ ECs to VEGF stimulation using the tube formation assay. Consistently, the ability of Casp-8^KD^ ECs to form capillary-like tube structures upon VEGF stimulation was also significantly attenuated (Suppl. Fig. 4B,C).

**Figure 3.**
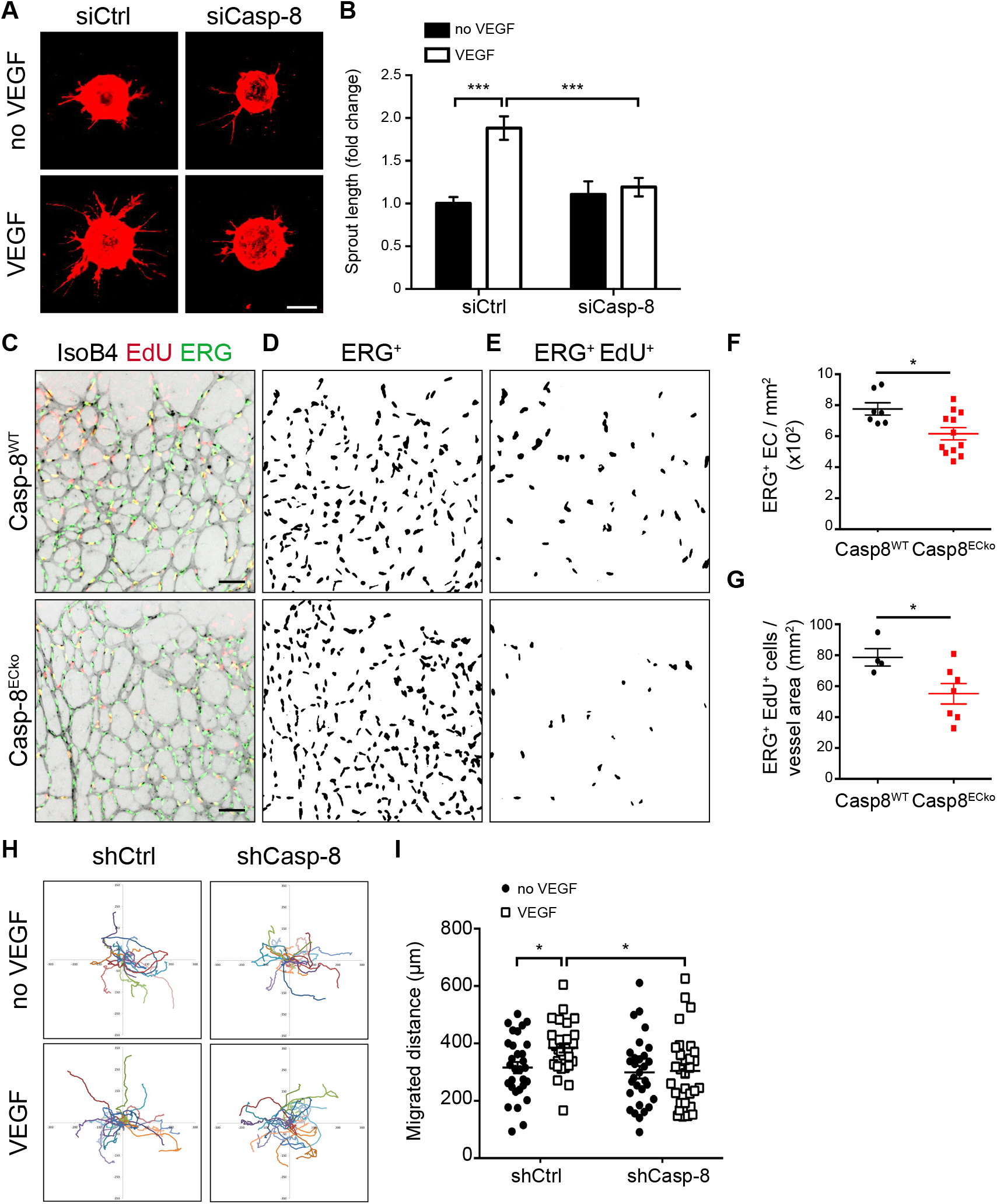
Loss of Casp-8 in ECs impairs sprouting, proliferation and migration. **A)** Representative images of the bead-sprouting assay using HUVECs transfected with control siRNA (siCtrl) or Casp-8 siRNA (siCasp-8) and treated with VEGF (50ng/ml) for 24h. **B)** Quantitative analysis of total sprout length showing that VEGF is not able to induce vessel sprouting in the absence of Casp-8. Approximately 20 beads per condition were quantified, n=4. **C)** Representative images of the retinal vasculature co-stained with IsoB4, EdU (labels proliferating cells) and ERG (labels ECs nuclei) in Casp-8^WT^ and Casp-8^ECko^ mice. IsoB4 single channel was transformed to grey colors and inverted with ImageJ for better visualization. **D,E)** Masks obtained by ImageJ of ERG^+^ cells (D) and of ERG^+^ EdU^+^ proliferating ECs (E). **F,G)** Quantification of total number of ECs per retina area (F; n=7 WT, 12 ECko) and proliferating ECs per vessel area (G; n=4 WT, 7 ECko), revealing lower absolute EC numbers and fewer proliferating ECs in Casp-8^ECko^ retinas compared to Casp-8^WT^ littermates. Data from 2 independent litters. **H)** Single cell-motility tracks of HUVECs infected with a control (shCtrl) or Casp-8 shRNA lentivirus (shCasp-8) and treated with VEGF (50ng/ml) for 12h. Migration origin of each cell was overlaid at the zero-crossing point. **I)** Quantification of the total migration distance of HUVECs in (H) showing that VEGF-induced migration was impaired in Casp-8^KD^ ECs. At least 30 cells per condition were quantified, n=3. For B,F,G,I data represent mean ± SEM (for B,I * *P* <0.05, *** *P* <0.001; two-way ANOVA with Bonferroni’s multiple comparisons test; for F,G * *P* <0.05; two-tailed unpaired Student *t*-test). Scale bars: A) 200μm; C) 50μm.

We next analyzed whether Casp-8 activity was required for the proper response of ECs to VEGF stimulation. Using a luminescent Glo-assay, we first confirmed that ECs have a basal Casp-8 activity, which could be blocked with the Casp-8 inhibitor Z-IETD-FMK (ZIETD, Suppl. Fig. 4D). As a positive control for our assay, we treated ECs with a combination of cycloheximide (CHX) and TNF to induce apoptosis, which strongly increased Casp-8 activity, as expected (Suppl. Fig. 4D). Functionally, blocking Casp-8 activity in ECs resulted in the same defects as the knockdown of Casp-8, as ZIETD treated ECs did not form sprouts upon VEGF stimulation (Suppl. Fig. 4E,F), indicating that Casp-8 activity was required for proper angiogenesis.

To assess if changes in EC proliferation could contribute to the reduced vascular area in Casp-8^ECko^ retinas, we performed proliferation experiments. ShRNA-mediated Casp-8 knockdown in HUVECs resulted in reduced proliferation upon stimulation with VEGF or FGF as determined by BrdU incorporation (Suppl. Fig. 4G). Also, siRNA transfected Casp-8^KD^ HUVECs had a reduced response to VEGF stimulation in the WST-1 cell proliferation and viability assay (Suppl. Fig. 4H). *In vivo*, the total number of ECs as determined by counting ERG+ nuclei per retina area (ERG is an EC specific transcription factor (34)) was reduced (Fig. 3C,D,F). EdU labeling of proliferating cells in P6 retinas showed significantly less EdU^+^/ERG^+^ cells per vessel area in Casp-8^ECko^ pups compared to Casp-8^WT^ littermates (Fig. 3C,E,G), indicating that reduced EC proliferation could account for the reduced vessel area.

EC migration is also required for proper expansion of the vascular network. We therefore checked whether knockdown of Casp-8 in ECs could also affect this process. In a classical scratch wound assay, VEGF stimulation of Casp-8^WT^ ECs led to almost 80% closure of the wound, whereas knockdown of Casp-8 reduced VEGF-induced wound closure (Suppl. Fig. 4I,J). To determine if Casp-8 affected migration independent of its effects on proliferation, we performed live imaging experiments and tracked the movement of non-dividing single Casp-8^KD^ ECs. Indeed, the total migration distance of ECs after 12h of VEGF stimulation was reduced when Casp-8 was knocked down (Fig. 3H,I).

Altogether, these results show that Casp-8 regulates EC proliferation and migration, which contributes to VEGF-induced sprouting.

### Loss of Casp-8 affects the organization of adherens and tight junctions in ECs

Apart of being important for vascular homeostasis and vessel stability, dynamic turnover of vascular endothelial cadherin (VE-cadherin) is crucial for proper vessel sprouting and elongation during angiogenesis (35–39).

As both sprouting and migration were impaired in Casp-8^ECko^ mice, and as Casp-8 has been shown to regulate the stability of cell junctions in the epidermis (40), we analyzed the distribution of VE-cadherin *in vivo* in the sprouting front and in the plexus of the growing vasculature of P6 retinas. By using an established image software analysis and classification key (35, 41), we distinguished between remodeling (‘active’) and stable (‘inhibited’) VE-cadherin patches. As expected, Casp-8^WT^ retinas presented a highly active VE-cadherin pattern in the sprouting front that was only slightly affected in Casp-8^ECko^ pups (Suppl Fig. 5A-C). More strikingly, Casp-8^ECko^ retinas had a destabilized, more discontinuous (‘active’) VE-cadherin staining in the back (plexus) of the retina, where Casp-8^WT^ retinas showed continuous (‘inhibited’) patches of VE-cadherin (Fig. 4A-C). Analysis of the distribution of Claudin-5 in the plexus, one of the main transmembrane proteins found in tight junctions, showed the same discontinuous pattern in the back of Casp-8^ECko^ retinas (Fig. 4A).

**Figure 4.**
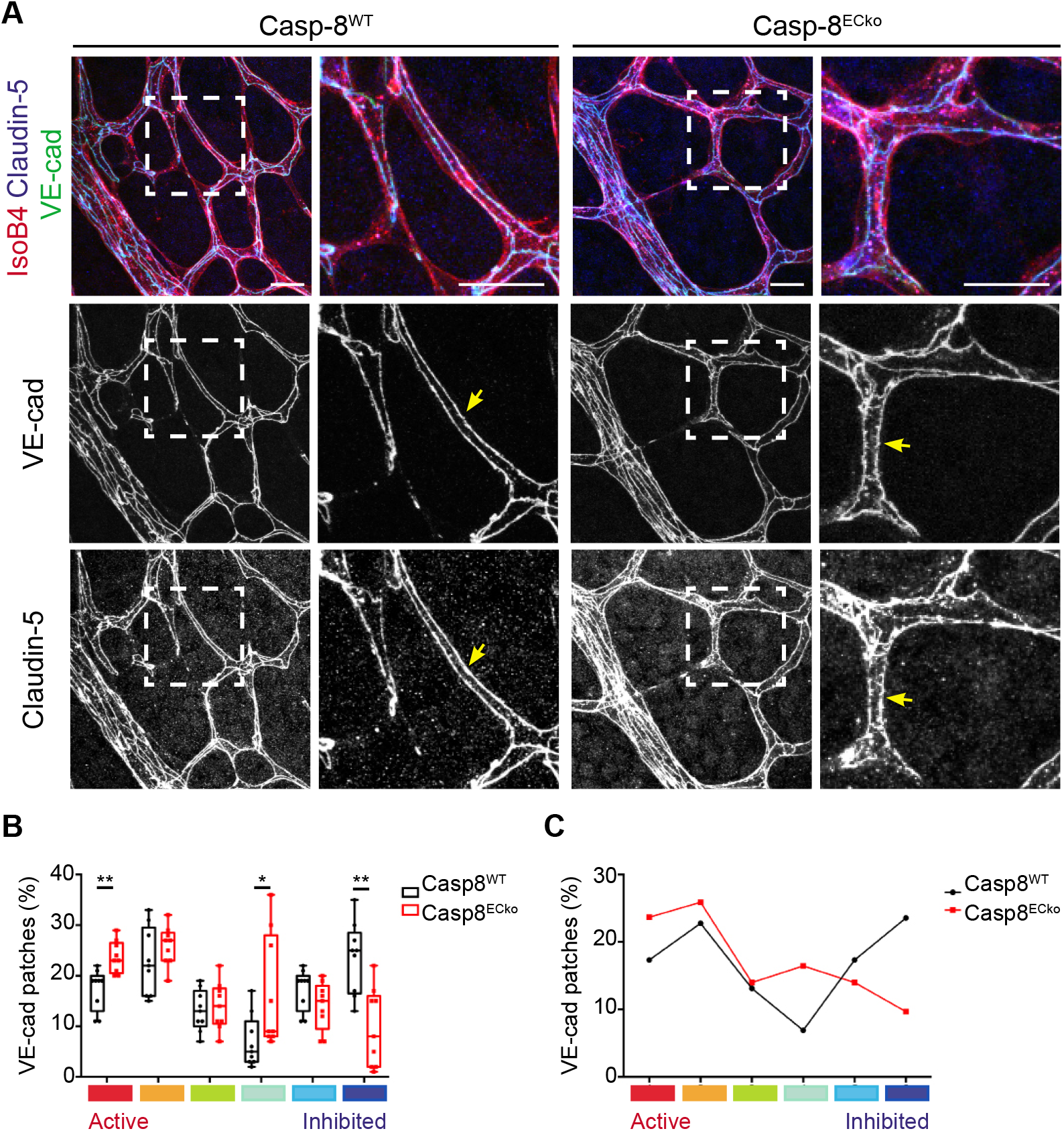
Casp-8 is necessary for maintaining EC junction stability at the retina plexus *in vivo*. **A)** Representative images of Casp-8^WT^ and Casp-8^ECko^ retinas stained with IsoB4, VE-cadherin and Claudin-5 (insets of left panels are shown on the adjacent right panels) showing that junctions are more serrated and discontinuous at the retina plexus in Casp-8^ECko^ compared to Casp-8^WT^ mice (yellow arrows). Images of VE-cadherin and Claudin-5 single channels were transformed to grey colors with ImageJ for better visualization. **B)** Quantification of the percentage of VE-cadherin patches showing a significant increase in the number of highly active VE-cadherin patches and a lower number of highly inhibited patches in Casp-8^ECko^ compared to Casp-8^WT^ mice. Each box shows the median percentage of patches of that type (line), and upper and lower quartiles (box). The whiskers extend to the most extreme data within 1.5 times the interquartile range of the box. (* *P* <0.05, ** *P* <0.01; Dirichlet regression model with two-tailed Mann Whitney test for each state; n=9 WT, 9 ECko). **C)** Average of the differential distribution of the percentage of VE-Cadherin patches in Casp-8^WT^ and Casp-8^ECko^ retinas. Scale bars: A) 20μm.

Next, we confirmed the VE-cadherin phenotype *in vitro*. Consistent with previous studies (42), confluent control shRNA transfected ECs (Casp-8^WT^ ECs) showed a continuous (‘inhibited’) VE-cadherin staining along the cell junctions (Fig. 5A), which became serrated with a rope-ladder pattern (‘active’) upon 30 or 60 min of VEGF stimulation (Fig. 5.B,C), as analyzed by measuring the average length of individual VE-cadherin patches (Fig.5D). In contrast, the distribution of VE-cadherin in confluent Casp-8^KD^ ECs was already discontinuous/serrated in unstimulated conditions and VE-cadherin did not further rearrange upon VEGF stimulation (Fig. 5A-D). Moreover, VE-cadherin at the cell perimeter were reduced at basal conditions (Fig. 5E). However, total VE-cadherin protein levels were unchanged (Fig. 5F), indicating that VE-cadherin was not degraded, but only mislocalized.

**Figure 5.**
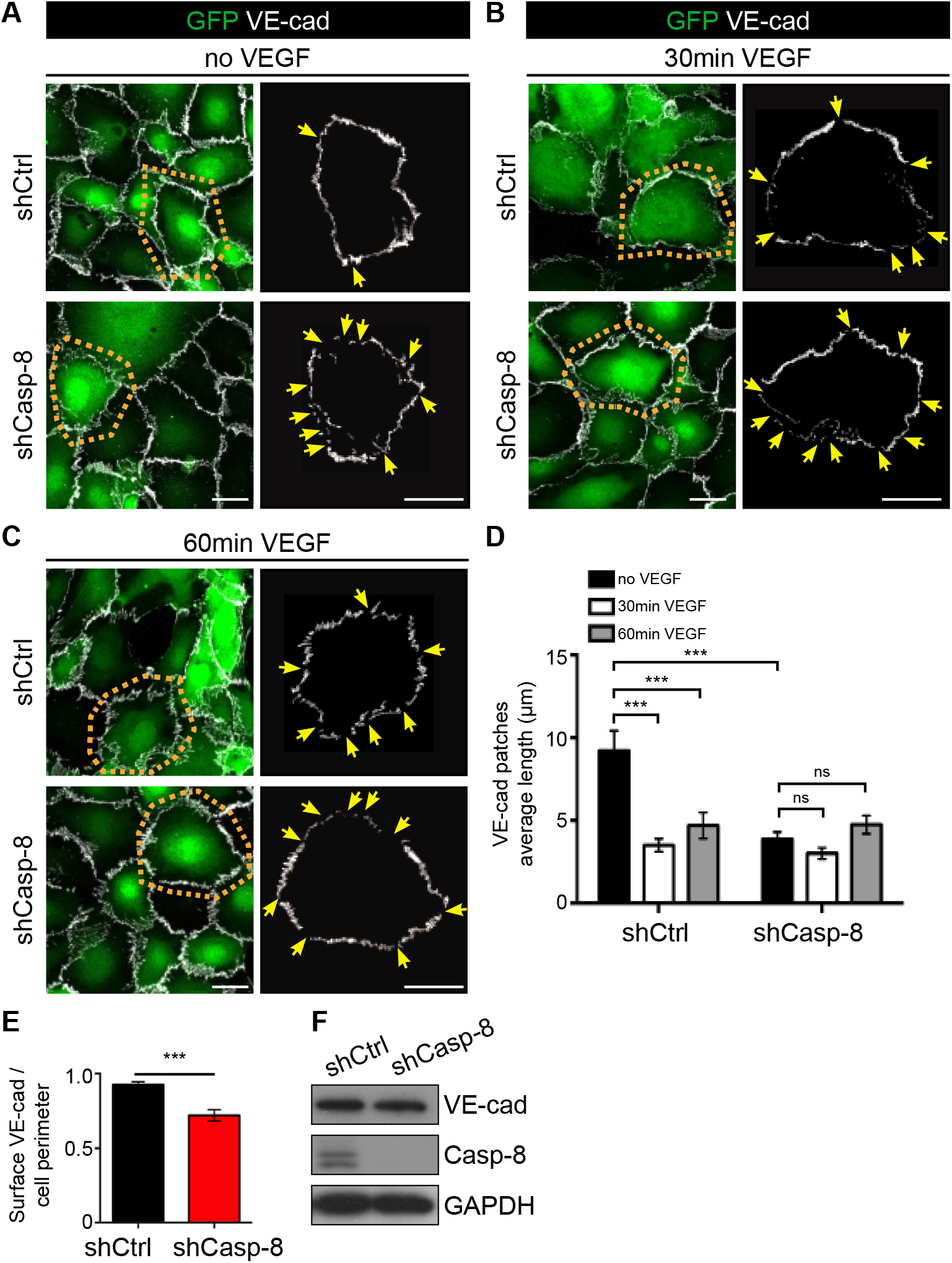
VE-cadherin distribution in ECs is affected in the absence of Casp-8 *in vitro*. **A-C)** Representative images of VE-cadherin staining in confluent HUVECs infected with shCtrl or shCasp-8 lentivirus (GFP^+^) with (B,C) or without (A) VEGF (50ng/ml) stimulation. Results of tracing VE-cadherin staining of the single cells within orange dotted insets is shown in the adjacent right panels. Yellow arrows point to empty VE-cadherin spots. **D)** Quantification of the average length of VE-cadherin patches showing that VEGF-induced VE-cadherin reorganization is impaired in Casp-8^KD^ ECs (*** *P* <0.001, ns: not significant; two-way ANOVA with Bonferroni’s multiple comparisons test). **E)** Quantification of the total amount of VE-cadherin per cell perimeter showing less VE-cadherin in Casp-8^KD^ ECs (*** P <0.001; two-tailed unpaired Student t-test).. **F)** Western blot showing unchanged total VE-cadherin protein levels in HUVEC infected with shCtrl or shCasp-8. For D,E at least 15 cells per condition were quantified, n=3. Data represent mean ± SEM. Scale bars: A-C) 20μm.

Taken together, our data shows that the absence of Casp-8 in ECs affects the proper formation of adherens and tight junctions in the postnatal retina, as well as the VEGF induced remodeling of VE-Cadherin *in vitro*.

### Loss of Casp-8 results in the basal activation of the p38 MAPK

We next pursued experiments aimed at identifying the signaling pathways involved in EC proliferation, sprouting and migration that could be altered in Casp-8^KD^ ECs. In the absence of Casp-8, phosphorylation of Akt, ERK and FAK was neither affected at basal conditions, nor upon VEGF stimulation (Suppl. Fig. 6A-D). However, knocking down Casp-8 or blocking its activity resulted in increased basal p38 MAPK phosphorylation (Fig. 6A-C). Still, Casp-8^KD^ ECs or HUVECs treated with ZIETD were able to further activate p38 upon VEGF stimulation (Suppl. Fig. 6A, E-G). As increased activation of p38 has been linked to the destabilization of VE-cadherin in the endothelium (43, 44), and as VE-cadherin distribution was already altered in basal conditions in Casp-8^KD^ ECs (Fig. 5A,D), we explored whether the increased basal levels of p-p38 in Casp-8^KD^ ECs were linked to the changes in the distribution of VE-cadherin. For this, we blocked p38 activity in confluent ECs with the specific p38 inhibitor SB203580 (45). Even though the length of individual surface VE-cadherin patches was only partially rescued (Fig. 6D,E), the total amount of VE-cadherin at the cell surface of Casp-8^KD^ ECs was fully recovered (Fig. 6F), indicating that indeed the basal activation of p38 was linked to VE-cadherin localization.

**Figure 6.**
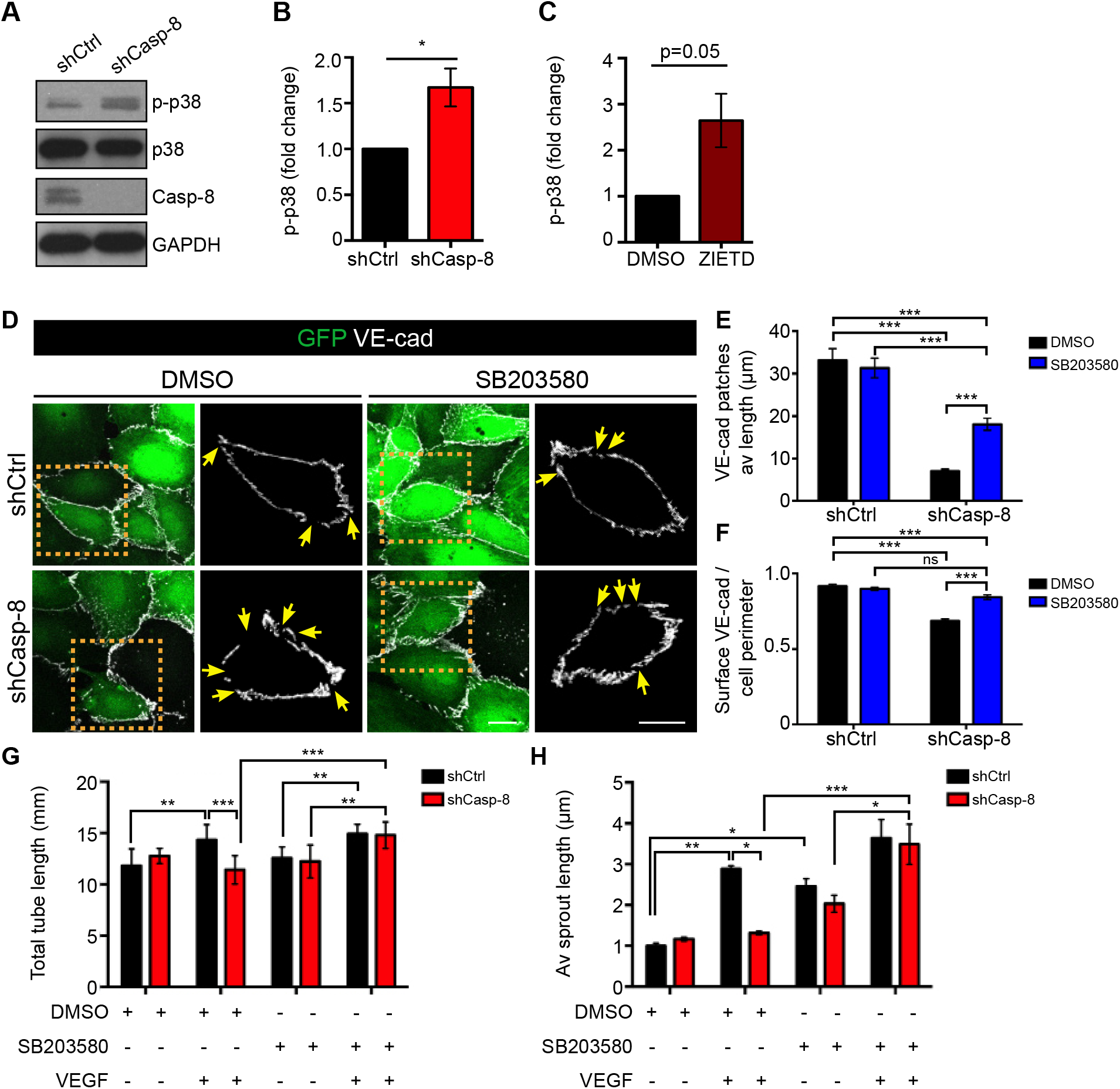
Loss of Casp-8 results in basal activation of p38 MAPK, VE-cadherin instability and defects in angiogenesis *in vitro*. **A)** Western blots showing increased phospho-p38 (p-p38) at basal conditions in Casp-8^KD^ (shCasp-8) ECs compared to control (shCtrl). **B)** Quantification of p-p38 as in (A), n=5. **C)** Quantification of p-p38 of HUVECs treated with ZIETD (10μM, 16h) at basal conditions, showing that blocking Casp-8 activity also induces increased basal p-p38, n=3. **D)** Images of VE-cadherin staining in shCtrl and shCasp-8 infected HUVECs treated with or without p38 inhibitor (SB203580, 1μM, 16h). Yellow arrows point to empty VE-cadherin spots. **E)** Quantification of VE-cadherin average patch length from cells as in (H). **F)** Quantification of the total amount of VE-cadherin per cell perimeter reveals that inhibition of p38 (SB203580) in Casp-8^KD^ ECs restores VE-cadherin to control levels. At least 15 cells per condition were quantified, n=3. **G)** Quantification of total tube length of HUVECs treated as in (G) showing that blocking p38 in Casp-8^KD^ ECs restores VEGF-induced tube formation. 3 fields per condition were quantified, n=4. **H)** Quantitative analysis of total sprout length showing that inhibition of p38 rescues VEGF-induced EC sprouting in Casp-8^KD^ ECs. Approximately 20 beads per condition were quantified, n=3. For B,D,F,G,I-L data represent mean ± SEM (for B,C * *P* <0.05; one sample *t*-test; for E-H * *P* <0.05, ** *P* <0.01, *** *P* <0.001, ns: not significant; two-way ANOVA with Bonferroni’s multiple comparisons test). Scale bars: D) 20μm.

To test whether the basal increase in p-p38 in Casp-8^KD^ ECs was also sufficient to functionally block VEGF-induced angiogenesis in Casp-8^KD^ ECs, we performed the tube formation and bead sprouting assays in the presence of the p38 inhibitor. Indeed, inhibition of p38 rescued the ability of Casp-8^KD^ ECs to respond to VEGF and to form both tubes (Fig. 6G and Suppl. Fig. 6H) and sprouts (Fig. 6H) of a similar length as Casp-8^WT^ ECs.

These results indicate that the loss of Casp-8 results in destabilization of EC junctions via an increased basal phosphorylation of p38. They also indicate that the increased basal p38 activity is responsible for the reduced response to VEGF, altogether resulting in overall impaired angiogenesis (see model in Fig. 7J).

**Figure 7.**
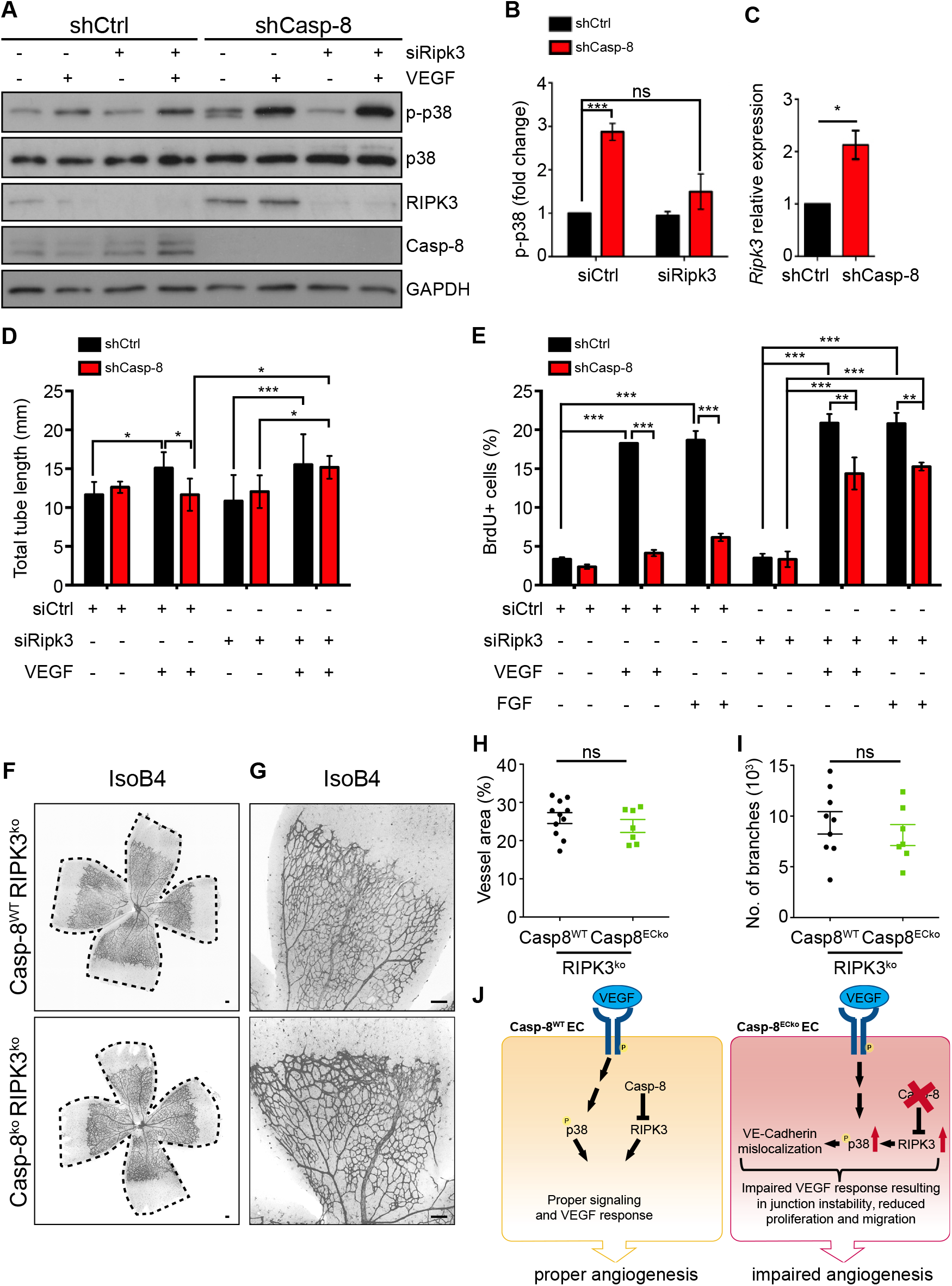
RIPK3 acts downstream of Casp-8 to regulate angiogenesis. **A)** Western blot showing that knocking down RIPK3 in Casp-8^KD^ ECs rescues the basal hyperphosphorylation of p38. Notice that shCasp-8 HUVECs have increased RIPK3 protein levels. **B)** Quantification of western blots at basal conditions as in (A), n=4. **C)** qPCR of shCtrl or shCasp-8 infected HUVECs showing increased mRNA expression of *Ripk3* in the absence of Casp-8, n=4. **D)** Quantification of total tube length of shCtrl or shCasp-8 infected HUVECs that were co-transfected with control siRNA or Ripk3 siRNA and treated with or without VEGF (50ng/ml) for 4h. 10 fields per condition were quantified, n=3. **E)** BrdU^+^ cells were quantified in control and Casp-8^KD^ ECs transfected with Ripk3 siRNA and with or without VEGF (50ng/ml) or FGF (50ng/ml) stimulation for 24h. Around 50 cells per condition were quantified, n=3. **F,G)** Representative images of the retinal vasculature stained with IsoB4 in Casp-8^WT^/RIPK3^ko^ and Casp-8^ECko^/RIPK3^ko^ mice. **H,I)** Quantitative analysis showing no differences on vessel area (H; n=11 WT, 7 ECko) and number of branches (I; n=9 WT, 7 ECko). Data from 4 independent litters. **J)** Working model summarizing the role of Casp-8 as a modulator of angiogenesis. Casp-8 inhibits RIPK3 to allow the proper response of ECs to VEGF stimulation (Casp-8^WT^). If Casp-8 is absent (Casp-8^ECko^), increased RIPK3 levels induce p38 hyperphosphorylation which in turn leads to an impaired response to VEGF stimulation and reduced angiogenesis. For B-D,E,H,I data represent mean ± SEM (for B,C * *P* <0.05, *** *P* <0.001, ns: not significant; one sample *t*-test; D,E * *P* <0.05, ** *P* <0.01, *** *P* <0.001; two-way ANOVA with Tukey’s multiple comparisons test; for H,I * *P* <0.05, ns: not signifcant; two-tailed unpaired Student *t*-test).

### Activation of p38 MAPK upon loss of Casp-8 is mediated by RIPK3

Casp-8 inhibits RIPK3 (12) and it has recently been shown that RIPK3 regulates vessel permeability via p38 in ECs (22). Therefore, we asked whether RIPK3 could act downstream of Casp-8 to regulate p38. Interestingly, we observed that Casp-8^KD^ ECs expressed higher RIPK3 protein (Fig. 7A) and mRNA levels (Fig. 7C), as also shown in epithelial Casp-8^KO^ mice (46). To investigate whether p38 was activated downstream of RIPK3 in Casp-8^KD^ ECs, we additionally knocked down RIPK3 and analyzed p38 phosphorylation. Knockdown of RIPK3 alone had no impact on either basal p-p38 nor on VEGF-induced p38 phosphorylation (Suppl. Fig 7A,B). However, the basal increase in p-p38 present in Casp-8^KD^ ECs was rescued to control levels in Casp-8^KD^/RIPK3^KD^ ECs (Fig. 7A,B), indicating that RIPK3 was responsible for the increased p-p38 levels observed in Casp-8^KD^ cells.

Consistent with these effects on p-p38, knockdown of RIPK3 also rescued the ability of Casp-8^KD^ ECs to respond to VEGF and to form tube-like structures to a similar extent as seen for Casp-8^WT^ and RIPK3^KD^ ECs (Fig. 7D, Suppl. Fig. 7C). In addition, proliferation induced by VEGF or FGF in Casp-8^KD^/RIPK3^KD^ ECs was also partially rescued compared to Casp-8^KD^ ECs, as measured by BrdU incorporation (Fig. 7E).

To confirm the role of RIPK3 in Casp-8^ECko^ mice *in vivo*, we crossed our Cdh5-(PAC)-CreERT2 x Casp-8^fl/fl^ mice with a RIPK3^ko^ mouse line (47). In line with our *in vitro* results, RIPK3 did not regulate angiogenesis under physiological conditions, as neither the vessel area nor the number of branchpoints (Suppl. Fig. 7D-G) in the retina of heterozygous or homozygous RIPK3^ko^ pups was affected at P6 compared to wildtype littermates. In contrast, the vascular defects in Casp-8^ECko^ mice were rescued in Casp-8^ECko^/RIPK3^ko^ mice as the vessel area (Fig. 7F-H) and number of branchpoints (Fig. 7I) were similar to Casp-8^WT^/RIPK3^ko^ mice.

In summary, these results show that in the absence of Casp-8, RIPK3 is mediating increased phosphorylation of p38 *in vitro* and impaired developmental vascular growth *in vivo*. Together, we propose that this RIPK3/p-p38 axis deregulates EC behavior resulting in impaired angiogenesis (Fig. 7J).

### Casp-8^ECko^ mice show reduced pathological angiogenesis in a model of oxygen-induced retinopathy

So far, our results indicate that Casp-8 is required for proper developmental angiogenesis and when absent, EC proliferation, migration and cell junction formation do not occur properly, resulting in reduced vessel sprouting and growth. To analyze the impact of these vascular defects in Casp-8^ECko^ mice at later developmental stages, we extended our tamoxifen treatment protocol to the first two postnatal weeks and analyzed Casp-8^ECko^ mice at P15 and P42 (Suppl. Fig. 8). At both stages, Casp-8^ECko^ mice did not show differences in the vessel area (Suppl. Fig.8B, E) or number of branches (Suppl. Fig. 8C,F) compared to control littermates. In addition, we analyzed vessel integrity by injecting 70kDa fluorescently labeled Dextran and could not detect any obvious difference in vessel permeability (Suppl. Fig. 8D, G) between genotypes. Taken together, this data suggests that compensatory mechanisms overcome the loss of Casp-8 in ECs.

Despite the better control of oxygen administration to preterm infants, ROP persists in extremely low gestational ages and birthweights and is still a clinical problem (48). ROP is characterized by two phases. In phase 1, exposure of preterm infants to high extra-uterine oxygen levels causes cessation of vessel growth. However, as the metabolic demand of the retina increases over time and as there is a lack of proper tissue oxygenation, upregulation of VEGF induces pathological neovascularization, consisting of extensive extra-retinal neovascular tufts in phase 2 of ROP (49). As our *in vitro* data showed that Casp-8^KD^ ECs were not able to respond to VEGF, we explored whether the loss of Casp-8 could also reduce pathological angiogenesis during ROP. We therefore applied the oxygen-induced retinopathy (OIR) model, which resembles very well the two phases of ROP (50, 51). In this model, high oxygen levels induce rapid vessel regression followed by upregulation of VEGF that causes pathological vessel sprouting and formation of abnormal and leaky vascular tufts.

For this, we exposed pups to high oxygen levels (75%) to induce vessel regression from P7 to P10 (Fig. 8A), as previously described (35). Subsequently, pups were returned to normoxia until P15, causing the now avascular retina to become hypoxic and upregulate VEGF, inducing the neovascular response. Cre recombination and Casp-8 deletion was induced on return to normoxia from P10 to P14 to evaluate the effect of Casp-8 loss on pathological neovascularization (Fig. 8A). Analysis of the retinal vasculature at P15 showed that pathological neovascularization (quantified by measuring the neovascular tuft area (52)) was strongly reduced in Casp-8^ECko^ compared to Casp-8^WT^ retinas (Fig. 8C, D), supporting the concept that blocking Casp-8 is beneficial to prevent disease progression. In addition to pathological neovascularization, normal angiogenesis gradually replaces the vessels lost from the central retina following exposure to high oxygen. Analysis of the avascular area showed a slight increase in Casp-8^ECko^ pups compared to wildtype littermates (Fig. 8B, E). This result is probably a combination of reduced pathological neovascularization and a reduced vessel regrowth, consistent with our finding that angiogenesis and the response to VEGF were reduced in Casp-8^KD^ ECs.

**Figure 8:**
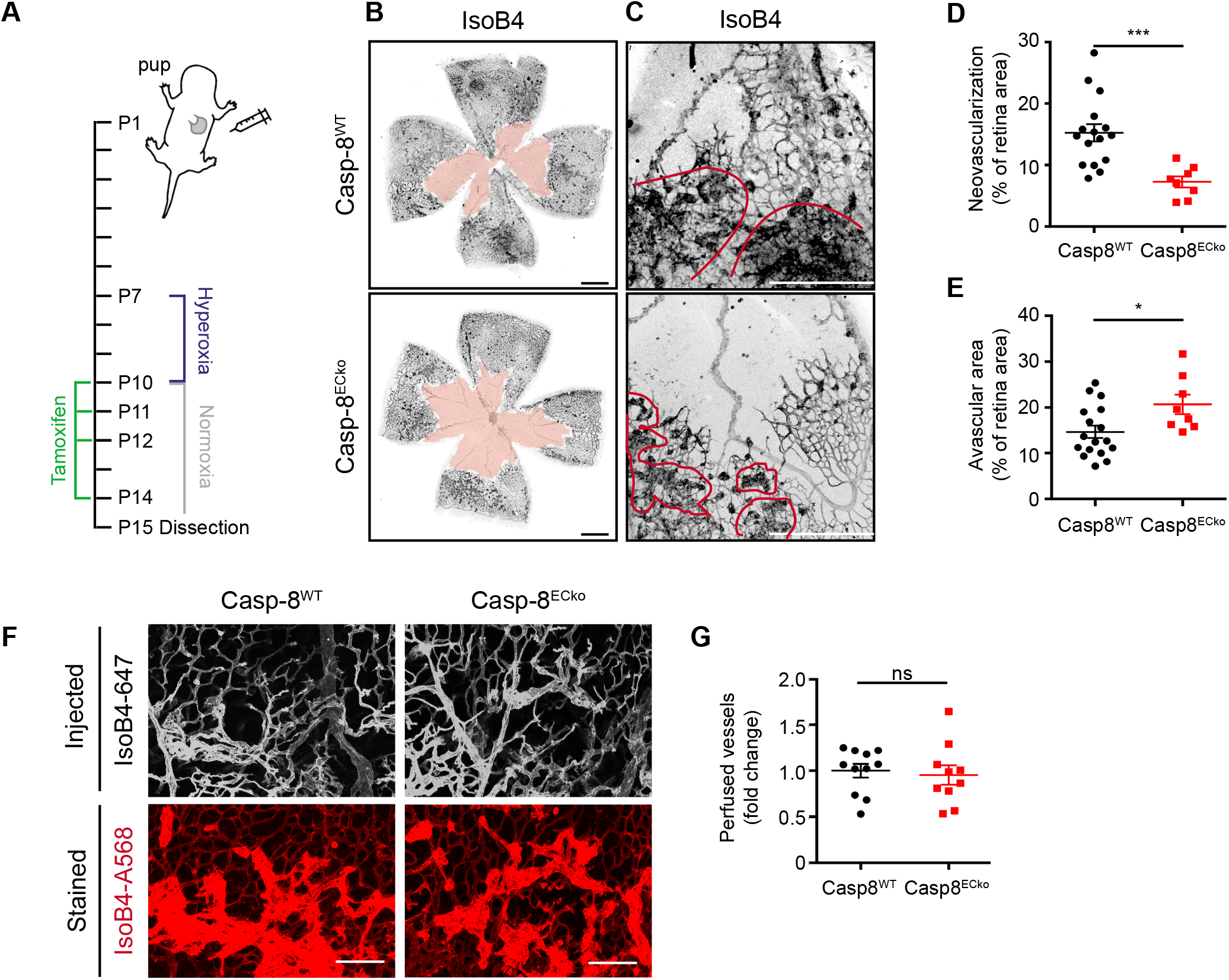
Loss of Casp-8 results in reduced neovascularization in the Oxygen Induced Retinopathy (OIR) model. **A)** Scheme showing the timeline of the used OIR protocol. Pups were placed in 75% oxygen (hyperoxia) from P7 to P10, then placed back to normal oxygen conditions (normoxia) until P15. Tamoxifen was injected at P10-12 and P14. **B,C)** Representative images of the retinal vasculature stained with IsoB4 in Casp-8^WT^ and Casp-8^ECko^ mice. Filled red space indicate the avascular area (B), red outlines indicate neovascular tufts (C). **D,E)** Pathological neovascularization is reduced in Casp-8^ECko^ mice (D) compared to Casp-8^WT^ littermates, while the avascular area was higher in Casp-8^ECko^ mice (n=16WT, 8ECko). **F,G)** Retro-orbital injection of IsoB4-647 did not reveal any difference in vessel perfusion among Casp-8^ECko^ and Casp-8^WT^ mice (n=11WT, 10ECko), as quantified by measuring Alexa647 fluorescent intensity in IsoB4-A568 labeled vessels. Data represent mean ± SEM from 3 independent litters (* *P* <0.05, ** *P* <0.01, *** *P* <0.001, ns: not significant; two-tailed unpaired Student *t*-test). Scale bars: B) 500μm; C) 250μm; F) 100μm.

Finally, to examine vessel integrity and functionality of the newly formed vasculature, we analyzed vessel perfusion by retro-orbital injection of IsolectinB4-A647 (which attaches to the vessel lumen and hence, labels perfused vessels), as previously described (53, 54). Then, the isolated retinas were costained with IsoB4-A568 to determine the vessel area. Analysis of IsolectinB4-Alexa647 per vessel area revealed efficient vessel perfusion in both genotypes (Figure 8F,H).

Taken together, targeting Casp-8 in ECs in a model of OIR, resembling ROP, could indeed be beneficial as it reduces the severity of tuft formation without compromising the newly formed vasculature.

## DISCUSSION

Components of both the intrinsic and extrinsic apoptosis signaling pathways have been described to be active in ECs and involved in different processes during the formation of the vascular system. While the intrinsic pathway of apoptosis (induced by the activation of Bax and Bak) is crucial for the regression and cell death of ECs from hyaloid vessels in the retina (55–58), this pathway controls EC number (8, 10, 59–61), but is not an active driver of vessel remodeling/pruning in the postnatal retina vasculature. On the other hand, activation of the extrinsic signaling machinery via activation of CD95 expressed in retinal ECs contributes to vessel development by regulating EC proliferation instead of cell death (62).

Casp-8 is a key protease in the extrinsic cell death signaling pathway that has both cell death-inducing and pro-survival functions (21). A basal Casp-8 activity is required for cell survival as it inhibits the activation of RIPK3 and thus the subsequent phosphorylation of MLKL leading to necroptosis (63). In pathological conditions, tumor-induced EC necroptosis has been described as a mechanism via which tumor cells extravasate and metastasize (23). However, in the same tumor model it has also been shown that RIPK3 regulates vessel permeability independent of EC necroptosis (22). Here we show that necroptosis is not involved in postnatal developmental angiogenesis, as the vascular defects in Casp-8^ECko^ mice are still present in Casp-8^ECko^/MLKL^ko^ retinas. Notably, proper angiogenesis was restored in Casp-8^ECko^/RIPK3^ko^ pups, indicating that the vascular impairments in Casp-8^ECko^ mice were mediated by RIPK3, thus assigning a necroptosis-independent function for RIPK3 in ECs during angiogenesis.

Cell death-independent functions of Casp-8 have been shown to be both activity dependent (16, 64) and independent (acting as a scaffold protein (14, 32)). While we do not rule out that Casp-8 could also act as a scaffold protein to regulate the angiogenic response of ECs, here we show that its activity is required. The cellular substrate for Casp-8 in ECs during angiogenesis has yet to be identified. However, as RIPK3 mRNA levels are upregulated in Casp-8^KD^ ECs, similar as in epithelial cells without Casp-8 (46), transcriptional regulators might be involved.

Mechanistically, Casp-8 deletion in ECs results in EC proliferation, sprouting and VE-cadherin subcellular distribution defects. Analysis of different signaling pathways revealed that Casp-8^KD^ ECs had increased basal activation of p38. Consistent with our findings, loss of Casp-8 in the epidermis has also been shown to induce the upregulation of p-p38 (65). Despite the fact that VEGF stimulation activated ERK, Akt, FAK and p38 in Casp-8^KD^ ECs, this did not result in a functional angiogenic response *in vitro*. However, when p38 activity was blocked, Casp-8^KD^ ECs responded normally to VEGF stimulation in the sprouting and tube formation assay. These findings indicate that the basal overactivation of p38 was already enough to functionally inhibit the response of Casp-8^KD^ ECs to VEGF and thus to impair angiogenesis. It was previously reported that p38 activity induces EC migration (66). However, in our experimental set-up, EC migration was not affected in Casp-8^KD^ ECs under basal conditions, even though p38 was already higher activated. As migration of Casp-8^KD^ ECs does not change in basal conditions, this suggests that p38-independent mechanisms might account for this phenotype. We could also link RIPK3 to p38 hyperactivation, as *in vitro* knockdown of RIPK3 in Casp-8^KD^ ECs rescued the basal phosphorylation of p38 to control levels. Further studies are needed to investigate how RIPK3 is regulating p38, and whether RIPK3 activity is required.

Specific loss of Casp-8 in the epidermis results in increased p38 phosphorylation (65) and loss of junction integrity in epithelial cells, independent of cell death (40). Here we show that loss of Casp-8 in ECs *in vitro* resulted in impaired distribution of VE-cadherin at the cell membrane, creating a serrated pattern. Moreover, Casp-8^KD^ ECs failed to induce VE-cadherin rearrangements required for proper angiogenesis upon VEGF stimulation. This phenotype was rescued when p38 was inhibited *in vitro*. So far, it is not very well understood how p38 regulates VE-Cadherin localization in physiological conditions. However, tumor transendothelial cell migration studies suggest that p38 does not act directly on VE-Cadherin, but rather promotes its internalization via the formation of stress fibers (67). We therefore hypothesize that the activation of p38 in the absence of Casp-8 could cause the VE-Cadherin defects via a similar mechanism.

*In vivo*, VE-cadherin stability was altered at the back of the retina of Casp-8^ECko^ EC pups, suggesting that EC-EC contacts fail to stabilize. Recent studies indicate that rather than its sole presence or absence (35–37, 39), it is the dynamic and heterogeneous expression of VE-cadherin in different areas of the developing vessel sprout that drives angiogenesis (38). Consistent with this current view, which suggests that EC junctions in the back of the retina have to be stabilized in order to allow proper sprout elongation in the front, we postulate that the increased number of vessel patches with serrated junctions in the back of Casp-8^ECko^ retinas blocks EC positional interchanges, ultimately resulting in a reduced number of sprouts at the vascular front and thus reduced angiogenesis.

In many pathological conditions such as ROP, proliferative diabetic retinopathy (PDR) and retina vein occlusion (RVO), among others, pathological retina neovascularization following tissue hypoxia is characterized by excessive extra-retinal neovascular tufts (50, 68). In fact, it is the aberrant neovascularization in these patients that causes the biggest clinical challenge (blindness), rather than vessel regression itself (68). Here, by applying a model of oxygen-induced retinopathy (OIR), resembling ROP and producing neovascular lesions resembling those of PDR or RVO, we describe that Casp-8 is mechanistically involved in the pathophysiology of ocular neovascular diseases, Indeed, Casp-8^ECko^ mice, were Casp-8 was deleted in ECs just during the neovascularization phase, presented reduced aberrant vascular tufts. Consistent with our findings that Casp-8^KD^ ECs have a reduced response to VEGF stimulation and thus show reduced VEGF-induced angiogenesis, we also observed a slight increase in the avascular area, suggesting that the newly regrowing vessels develop slower in Casp-8^ECko^ retinas compared to Casp-8^WT^ ones (however, we cannot exclude that in this specific setting, necroptosis contributes to the increased avascular area observed in Casp-8^ECko^ mice). Nevertheless, based on the fact that in physiological conditions Casp-8^ECko^ retinas finally achieve a normal and functional vasculature, we propose that inactivation of Casp-8 in an ischemic retinopathy pathological setting, such as ROP, is beneficial as it will inhibit aberrant neovascularization, and, in a similar way as in development, the avascular area will be finally recovered.

## EXPERIMENTAL PROCEDURES

### Mouse lines and treatment

Casp-8^fl/fl^ mice (69) were crossed with Cdh5(PAC)-CreERT2 mice (24), to specifically delete Casp-8 in the the endothelium upon tamoxifen treatment. MLKL^ko^ mice were provided by M. Pasparakis (CECAD, Cologne). RIPK3^ko^ mice (47) were provided by Genentech (South San Francisco, CA, USA). Pregnant females were intraperitoneally injected with 2mg tamoxifen/30g body weight from E8.5 until E10.5. Embryos were dissected at E13.5 and assessed for heart beat (alive), death or presence of malformations such as bleedings or (cardiac) edema (Suppl. Fig. 1B). Recombination in pups was induced by intragastric injection of 50μl tamoxifen (1mg/ml) at postnatal day (P)1–P3 and P5 as described in Figure 1A. For analysis at P15 and P42, pups received additional tamoxifen injections (i.p.) at P8, P11 and P14.

### Oxygen induced retinopathy model

To induce retinal pathological angiogenesis in Casp-8^ECko^ mice, the OIR model was used as previously described (35). From P7 to P10, pups were placed in 75% oxygen (hyperoxia); at P10 pups were returned to normal oxygen conditions (normoxia) until P15 (35). Cre recombination was induced by i.p. injection of 50μl tamoxifen (2mg/ml) at P10 – P12 and P14. To analyze vessel perfusion, pups at P15 were retro-orbitally injected with IsoB4-Alexa 647 (5mg/kg body weight) as previously described (54) and sacrificed after 5 min. Retinas were isolated and fixed in 4%PFA/PBS for 1h at room temperature (RT). Conditions used for the retina staining are specified in the “Retina staining” section. Quantification of the size of the avascular area and neovascular tufts was performed in Adobe Photoshop blind to experimental conditions as previously described (52). To quantify vessel perfusion, mean fluorescent intensity of injected IsoB4-Alexa 647 was normalized to the intensity of IsoB4-Alexa 568 stained vasculature as previously described (54).

### Analysis of EC proliferation *in vivo*

EdU (Thermo Scientific) was injected at a concentration of 100 μg/g body weight into P6 Casp-8^ECko^ pups as previously described (70) 2.5 h before culling. Eyes were fixed in 4%PFA for 1.5h at 4°C. EdU^+^ cells in the retina were detected with the Click-iT^™^ EdU Alexa Fluor^™^ 488 Imaging Kit (Thermo Scientific) according to manufacturer’s instructions. ECs were counterstained with ERG and IsoB4. Eight pictures/pup of the angiogenic front were imaged at 20x magnification with a LSM 510 META. Quantification was done blind to experimental conditions. The number of proliferating ECs is expressed as EdU^+^/ERG^+^ cells normalized to the IsoB4^+^ vessel area per field of view.

### Whole mount staining of the yolk sac vasculature

Yolk sacs were fixed overnight in 4% PFA/PBS at 4 °C. After permeabilization in 1% Triton X-100/PBS for 1h at RT and blocking in 0.2% BSA, 5% normal donkey serum (Dianova), 0.3% TritonX-100 in PBS for 2h at RT, primary antibody Endoglin (1:200, MAB1320, R&D Systems) was incubated overnight at 4 °C in blocking solution. The appropriate Alexa Fluor^™^-conjugated secondary antibody was incubated overnight at 4°C. Whole mount images were acquired on a LSM 510 META confocal microscope.

### Retina dissection, processing and staining

Eyes were collected and enucleated in PBS. For VE-Cadherin stainings, eyes were fixed in 2% PFA/PBS for 1h at 4°C. For all other stainings, eyes were fixed for 1.5h in 4% PFA/PBS at 4°C. Retinas were dissected, permeabilized with PBS containing 1% BSA, 0.3% Triton X-100 for 1h and incubated with primary antibodies at 4°C overnight. Antibodies used were Alexa Fluor^™^ 594-conjugated IsolectinGS-IB4 (IsoB4) (1:250, ref.I21413, Thermo Scientific), Casp-8 (1:100, ref.4927, Cell Signaling), ERG (1:200, ab92513, Abcam), Collagen IV (1:200, CO20451, Biozol), Desmin (1:200, ab15200, Abcam), VE-cadherin (1:100, ref.555289, BD Bioscience), Claudin-5 (1:100, ref.34-1600, Thermo Scientific), cCasp-3 (1:100, ref.661, Cell Signaling). After washing with PBS, retinas were incubated for 2h at RT with the respective secondary antibody. TUNEL assay (Roche) was performed according to manufacturer’s instructions. Briefly, retinas were fixed for 2h in 4% PFA/PBS at RT. Permeabilization was done in 0.5% Triton X-100/0.1% citrate in PBS for 3h at RT. Then, labelling of TUNEL^+^ cells was done for 1h at 37 °C. After washing, samples were stained with FITC-conjugated IsoB4 (1:250, ALX-650-001F-MC05, Enzo Life Sciences) in 1% BSA, 0.3% Triton X-100/PBS for 1h at RT. Retinas were flat-mounted and analyzed using a confocal fluorescence microscope (LSM 510 META or LSM800; Carl Zeiss).

### Analysis of VE-cadherin localization in ECs *in vivo*

For analysis of VE-Cadherin in the retina of P6 pups, high magnification pictures (63x) were acquired. Two pictures from either the angiogenic front or the back of the retina were randomly taken in areas adjacent either to a vein or an artery. Representative inset pictures show the most active or inhibited pattern of VE-cadherin in Casp-8^WT^ and Casp-8^ECko^ pups. Quantification of the percentage of VE-cadherin patches was done blind to experimental conditions in MATLAB as previously described (35).

### *In situ* hybridization in the retina

*In situ* hybridization (ISH) was performed with minor modifications as previously described (71). In brief, eyes of P6 pups were removed and fixed for 20 min in 4% PFA on ice. After isolation and dissection, retinas were stored in methanol at −20°C overnight. Hybridization was performed at 66°C. Primer sequences to generate probes to detect mouse Casp-8 are: Casp-8-Fwd 3’-TTTCCACATCAGTCGGTGGG-5’ and Casp-8-Rev 3’-CTCTTGGCGAGTCACACAGT-5’. After color development, retinas were extensively washed with PBS and counterstained with Alexa Fluor^™^ 594-conjugated IsoB4. Bright field and fluorescent images were acquired simultaneously using a Zeiss LSM 800 (20x objective). Bright field images were processed and background corrected with a rolling ball algorithm in NIH ImageJ.

### Lung endothelial cell isolation

Murine lung ECs were isolated by MACS^®^ Technology (Miltenyi Biotec). Lungs of P6 mouse pups were isolated and kept in ice-cold HBSS containing 10% FCS. Tissue from Casp-8^WT^ and Casp-8^ECko^ pups from the same litter were pooled and digested in collagenase I/DNAse I (0.1mg/ml, Roche) at 37°C. The digested tissue was filtered, incubated with Red Blood Cell Lysis Buffer (Roche) and washed several times with PEB buffer (0,5% BSA, 2 mM EDTA in PBS). Negative selection of CD45^+^ cells was performed by incubation of the cell suspension with CD45 MicroBeads (1:10) for 15min at 4°C. Cell suspensions were applied onto LS columns and unlabeled cells were collected on ice. Enrichment of ECs was performed using CD31 MicroBeads (1:10) in PEB buffer for 15 min at 4°C and application on MS columns. The positive cell fraction was eluted from the columns with PEB buffer and directly frozen in RLT buffer (Qiagen) for RNA isolation.

### Cell culture

Human umbilical vein endothelial cells (HUVECs; PromoCell) were cultured in Endopan 3 Kit for endothelial cells (PAN-Biotech) supplemented with 10% FBS, 100 U/ml penicillin and 100 μg/ml streptomycin (both Gibco^®^ by Life Technologies) in a 5% CO_2_ humidified incubator at 37°C. Cells from passages 2 to 5 were used for the experiments. When indicated, cells were starved in growth factor-free Endopan 3 supplemented with 2% FBS, 100 U/ml penicillin and 100 μg/ml streptomycin.

### RNA extraction and quantitative Real-Time PCR analysis

RNA was extracted with the mini RNA extraction kit (Qiagen). RNA samples were transcribed to complementary DNA using Maxima Reverse Transcriptase (Thermo Scientific) or SuperScript VILO cDNA Synthesis Kit (Invitrogen). qRT-PCR was performed using Fast SYBR Green Master Mix (Thermo Scientific). Either *GAPDH* or *ActB* were used as housekeeping genes. All qPCR results from cultured cells were obtained from at least 3 independent experiments. Results from lung isolated ECs are from 4 independent litters.

### siRNA transfection

HUVECs were seeded in a 6 well plate (9×10^4^−11×10^4^ cells/well). The next day, cells were transfected with MISSION^®^ siRNA Universal Negative Control (Sigma-Aldrich) or previously validated siRNAs for Casp-8 (72) and/or RIPK3 (22) using Oligofectamin (Life Technologies), according to the manufacturer’s instructions. The final concentration of siRNA solution was 200nM.

### Lentiviral vectors and virus production

For silencing experiments, previously used and validated lentiviral vectors to express shRNAs against Caspase-8, c-FLIP or control (29, 73, 74) were used. Lentiviruses were produced by transfection of HEK293-T cells with the calcium phosphate method with the corresponding vectors. Lentivirus-containing supernatants were collected 48/72 h after transfection and concentrated by ultracentrifugation at 22,000 rpm for 90 minutes at 4 °C.

### Analysis of VE-cadherin localization in ECs *in vitro*

For VE-cadherin staining in HUVECs, 1×10^5^ infected cells (shCtrl or shCasp-8) were seeded and allowed to attach overnight. Cells were starved overnight and stimulated with 50ng/ml VEGF for the indicated time points. After fixation in 4% PFA/PBS for 30 min at RT and permeabilization in 0.5% Triton X-100/PBS for 10 min, blocking in 1%BSA, 20% normal donkey serum (Dianova) in 0.2% Triton X-100/PBS was done for 1h at RT. Samples were incubated with primary antibody (anti-VE-cadherin, 1:200, 610252, BD Bioscience) for 2h at RT followed by incubation with the appropriate Alexa Fluor^™^-conjugated secondary antibody for 2h at RT. Nuclei were counterstained with DAPI (1:1000, D1306, Invitrogen). After mounting, random fields of view were imaged with a Zeiss LSM 800 (63x objective). VE-cadherin at the cell perimeter of single, non-adjacent cells was manually traced with ImageJ. Total VE-cadherin at the cell perimeter was calculated as the sum of all individual VE-cadherin patches. At least 15 cells per condition were quantified blind to the experimental condition from 3 independent experiments.

### *In vitro* BrdU incorporation

To analyze EC proliferation *in vitro*, 25×10^3^ infected HUVECs (shCtrl or shCasp-8) were plated in 0.1% gelatin/water coated coverslips. After overnight starvation, cells were treated with or without VEGF (50ngml) or FGF (50n/ml), for 24 h. BrdU (10μM) was added and incubation of the cultures continued for 4h at 37°C. Cells were fixed in 4% PFA/PBS for 20min and permeabilized and blocked in 2% BSA 0.3% Triton X-100/PBS for 30min at RT. After unmasking with ice-cold HCl, neutralization with sodium borate buffer (0.1M Na2B4O7 in water, pH 8.5) for 15min was done prior to primary antibody incubation. An anti-BrdU antibody (1:250, OBT0030, Oxford Biotechnology) was incubated in blocking solution overnight at 4 °C and an appropriate Alexa Fluor^™^-conjugated secondary antibody was incubated for 2h at RT. Nuclei were counterstained with DAPI (1:1000, D1306, Invitrogen). Cells were mounted and imaged in a wide field microscope (Zeiss Axiovert 200) equipped with an Axiocam MRm camera (40x objective). Around 100 cells per condition of 3 independent experiments were quantified blind to experimental conditions.

### WST-1 assay

The WST-1 cell proliferation and viability assay (Sigma-Aldrich) was done according to the manual as previously described (33). Briefly, transfected HUVECs were plated in a 96 well plate and allowed to attach overnight. Then, cells were starved overnight and stimulated with VEGF (50ng/ml) or control vehicle for 24h.

### Fibrin gel bead sprouting assay

Fibrin gel bead sprouting assay was performed as previously described (75). Briefly, cytodex 3 microcarrier beads (GE Healthcare) were coated with siRNA-transfected or shRNA-infected HUVECs (mixed at 200 cells per bead), and embedded in 2 mg/mL fibrin gels (2 mg/mL fibrinogen (Calbiochem), 0.625 Units/mL thrombin (Sigma-Aldrich) and 0.15 Units/mL aprotinin (Sigma-Aldrich)). Beads were cultured in 2% FBS growth factor-free Endopan 3 medium, in presence or absence of 50ng/ml VEGF. After 24 hours the culture was fixed with 4% PFA for 15min, blocked in 1% BSA, 0.2% TritonX-100-PBS and incubated with 1μg/mL Phalloidin-fluo (94507, Sigma-Aldrich) for 2h. Confocal images were taken with a Zeiss LSM 510 microscope and analyzed by NIH ImageJ. Quantification was done blind to experimental conditions. Approximately 20 beads per condition were quantified from 4 independent experiments.

### Single endothelial cell tracings

Single EC tracings were performed as previously described (33). Infected ECs with shRNA control or shCasp-8 were plated on 6 well plates and stimulated with 50ng/ml VEGF or control vehicle (after overnight starvation). Phase contrast images of 10 fields of view per well were acquired every 10 min over the course of 12h using an inverted NikonTi microscope (with a Nikon Plan Fluor 10x NA 0.3 objective) equipped with an environmental box from Oko Lab for temperature, CO_2_ and humidity control. EC migration was traced automatically using NIS Elements 4.5 software. At least 30 cells per condition were quantified blind to experimental conditions from 3 independent experiments.

### Tube formation

Tube formation assays were performed in μ-Slide Angiogenesis wells (ibidi GmbH, Germany) as previously described (33, 76) using 10 μl of growth factor-reduced Matrigel (BD Bioscience) per well. In brief, siRNA transfected or lentivirus infected HUVECs resuspended in 2% FBS growth factor-free Endopan 3 medium, with or without VEGF (50ng/ml), were seeded onto the polymerized matrigel. Wherever indicated, prior to seeding on Matrigel, cells were pre-treated overnight with SB203580 (1μM). Cells were incubated for 4h in a humidified chamber at 37°C, 5% CO2 before analysis. Images were acquired with a wide field microscope (Zeiss Axiovert 200) equipped with an Axiocam MRm camera and a 5x objective. At least 3 fields of view per condition were acquired. Quantification of three independent experiments was done blind to experimental conditions with the Angiogenesis analyzer tool of NIH ImageJ.

### Scratch assay

siRNA transfected HUVECs were seeded in a 6 well plate. When cells were confluent, they were starved in 2%FBS growth factor-free Endopan 3 medium overnight. A wound was induced by scraping the cell monolayer with a P200 pipet tip and cells were stimulated with or without VEGF (50ng/ml). A picture was acquired at time-point zero and 12h after incubation at 37°C.The percentage of wound closure between 0h and 12h of VEGF stimulation was analyzed with NIH ImageJ, blind to experimental conditions. Results are from at least 3 independent experiments. For each treatment 8-10 fields of view were analyzed.

### *In vitro* cell death assay

Infected HUVECs (1×10^5^ cells/well in 6 well plates) were treated with TNF-α (100ng/ml, Preprotech) and TRAIL (100ng/ml, homemade) in normoxia or hypoxia (0,1% O2, BioSpherix, X2, Exvivo System) during 24h. After treatment, cells were washed with PBS, and stained with PI (40ug/ml) while treating with RNase (100ug/ml). Quantitative analysis of PI^+^ cells was carried out in a FACSCalibur cytometer using the Cell Quest software (Becton Dickinson).

### Caspase-8 Glo assay

The Caspase-8 Glo^®^ assay was performed in combination with the CellTiter-Fluor^™^ Cell Viability Assay (both from Promega) according to the manual. In brief, 6×10^3^ HUVECs per well were seeded in a 96-well plate. After overnight starvation, cells were treated with Z-IETD-FMK (10μM) or TNF (100ng/ml) + CHX (10μg/ml) for 4h before cell viability and Casp-8 activity were measured. Data is expressed as relative units (RU) resulting from the Caspase-8-Glo [RLU]/ Cell Viability [RFU] ratio.

### Immunoblotting

HUVECs were starved overnight and stimulated with VEGF (50ng/ml). When indicated, cells were additionally treated with ZIETD (10μM) in starvation medium overnight. Cells were lysed in lysis buffer (20mM Tris, 137mM NaCl, 2mM EDTA, 10% glycerol, Roche Protease and Phosphatase Inhibitor Cocktail). Proteins were separated by SDS-PAGE and immunoblotted according to standard protocols. GAPDH (1:10000, SC-47724, Santa Cruz), p-p38 (1:1000, ref.36-8500, Invitrogen), p38 (1:1000, ref.9228, Cell Signaling), p-Akt Ser473 (1:1000, ref.4060, Cell Signaling), Akt (1:1000, ref.9272, Cell Signaling), p-FAK Tyr397 (1:1000, ref.3283, Cell Signaling), FAK (1:1000, ref.3285, Cell Signaling), p-ERK p44/42 (1:1000, ref.9106, Cell Signaling), ERK (1:1000, ref.9102, Cell Signaling), RIPK3 (1:1000, ref.13526, Cell Signaling), Casp-8 (1:1000, ALX-804-429, Enzo), c-FLIP (1:1000, AG-20B-0056, AdipoGen).

### Statistics

Results are expressed as mean ± SEM. To calculate statistical significance, the two tailed Student’s t-test, one sample t-test, one-way ANOVA or two-way ANOVA followed by Bonferroni’s multiple comparisons test were used and indicated in the appropriate figure legend. To calculate statistical significance in Fig. 4 (analysis of VE-Cadherin distribution *in vivo*), Dirichlet regression model was applied for the analysis of binned data. Additional Mann-Whitney tests for each state are reported to indicate states with strong differences between groups. All calculations were performed using Prism software.

### Study approval

All animal procedures were conducted in accordance with European, national, and institutional guidelines. Protocols were approved by local government authorities (Regierungspräsidium Karlsruhe, Germany).

## Author contributions

Conceptualization: N.T., A.F.V., R.Y., C.R.A.; Methodology: N.T., A.F.V., R.Y., C.R.A.; Investigation: N.T., A.F.V., R.Y., I.P., S.L.P., X.W., R.M.P., L.C., W.W.W., L.C., B.S., T.H., C.M.,P.H., C.R.A.; Resources: C.R.A., H.G.A., A.L.R.; Writing - Original draft: N.T., A.F.V., R.Y., T.S., C.R.A.; Writing – Review & Editing: N.T., A.F.V., R.Y., I.P., S.L.P., X.W., R.M.P., W.W.W., L.C., B.S., H.J.G., L.C.W., M.M., A.L.R., T.S., H.G.A., T.H., C.M.,P.H., C.R.A.; Supervision: C.R.A., A.L.R., T.S., H.G.A.; Project Administration: C.R.A.; Funding Acquisition: C.R.A.

## Acknowledgements

We thank Manolis Pasparakis, Stephen Hedrick and the UCSD as well Bruno Köhler and Genentech for providing MLKL^ko^, Casp-8^flox/flox^ and RIPK3^ko^ mice, respectively. We thank Katie Bentley for her help with the MATLAB image analysis for VE-Cadherin stainings *in vivo*. We thank the Nikon imaging Center of the University of Heidelberg for their support. We thank Heike Adler and Melanie Richter for technical assistant and the Ruiz de Almodovar lab for useful discussions. N.T. was supported by an HBIGS PhD fellowship; I.P. was supported by Becas Chile. X.W. was supported by an Alexander Von Humboldt postdoctoral fellowship. T.S. is supported by DFG SCHM 2560/3-1. CRA is supported by DFG grant RU 1990/1-1, ERC (ERC-StG-311367) and the Deutsche Forschungsgemeinschaft (DFG)-SFB873; FOR2325 and SFB1366 (Project number 394046768-SFB 1366). L.C. is supported by an NHMRC Project Grant (1125536) and the L.E.W Carty Charitable Fund.

---

**Suppl Fig 1. Knockout of Casp-8 in ECs is lethal during embryonic, but not postnatal development.**

**A)** Casp-8^WT^ and Casp-8^ECko^ mouse embryos at E13.5. Upper and middle panels: embryo overview showing developmental arrest or severe haemorrhages (white arrows) in Casp-8^ECko^ embryos. Lower panels: immunostaining for the EC marker Endoglin, showing impaired yolk sac vasculature in Casp-8^ECko^ embryos (E13.5). **B)** Table of embryonic lethality of Casp-8^ECko^ embryos at E13.5. **C)** qRT-PCR of isolated lung ECs showing purity of the CD31^+^ fraction. Values were normalized to the CD31^−^ fraction. **D)** qRT-PCR of isolated lung ECs showing knockout efficiency in Casp-8^ECko^ pups (results from 3 independent litters; ** *P* <0.01; one-sample *t*-test). **E)** No differences between genotypes were observed on mouse weight (n=25 WT, 22 ECko). **F)** Representative images from Casp-8^WT^ and Casp-8^ECko^ mice. **G)** No differences between genotypes were observed on survival (n=25 WT, 22 ECko). **H)** *In situ* hybridization (mRNA) or immunofluorescence (protein) showing Casp-8 expression at the front and the back (plexus) of P6 mouse retinas. Data represent mean ± SEM. Scale bars: A) 1cm (upper panel) and 100μm (lower panel); F) 5mm; H) 100μm.

**Suppl Fig. 2. Knockdown of Casp-8 in ECs does not increase cell death *in vitro*.**

**A)** Western blot showing protein knockdown efficiency after infecting HUVECs with a lentivirus carrying shRNAs for c-FLIP or Casp-8. **B,C)** Quantification of apoptotic/necrotic HUVECs (PI^+^) in normoxia (B) or hypoxia (C) showing that knockdown of c-FLIP (shFLIP), after the indicated treatments, increases cell death while knockdown of Casp-8 (shCasp-8) does not, n=3. Data represent mean ± SEM (** *P* <0.01, *** *P* <0.001, ns: not significant; two-way ANOVA with Bonferroni’s multiple comparisons test).

**Suppl Fig. 3. MLKL knockout pups do not present impaired postnatal angiogenesis.**

**A,B)** Representative images of whole mount P6 retinas of MLKL^WT^, MLKL heterozygous (MLKL^Het^) and MLKL^ko^ mice stained with IsoB4 (ECs) (A) and higher magnifications of the retina front (B). Black dashed lines highlight the total retina area. **C,D)** Quantification of vessel area (C; n=4 WT, 6 Het, 4 ko) and number of branches (D; n=3 WT, 4 Het, 4 ko). For C,D data represent mean ± SEM from 2 independent litters (ns: not significant; one-way ANOVA with Bonferroni’s multiple comparisons test). Scale bars: A) 100μm; B) 50μm.

**Suppl Fig. 4. Knockdown of Casp-8 impairs VEGF-induced tube formation, EC proliferation and migration *in vitro*.**

**A)** Western blot showing the knockdown efficiency of the Casp-8 siRNA. **B)** Bright field representative images of the tube formation assay. HUVECs were transfected with control siRNA (siCtrl) or Casp-8 siRNA (siCasp-8) and treated with VEGF (50ng/ml) for 4h. **C)** Quantification of the total tube length as in (B) showing that Casp-8 knockdown inhibits VEGF-induced tube formation. 5 fields per condition were quantified, n=3. **D)** Casp-8 activity was measured with a Casp-8 Glo kit. HUVECs’ basal Casp-8 activity can be blocked with the Casp-8 inhibitor ZIETD (10μM) or induced by cycloheximide (CHX, 1μg/ml) plus TNF (100ng/ml) for 4h, n=3. Data represent mean ± SEM (** *P* <0.01, *** *P* <0.001; two-tailed unpaired Student *t*-test). **E)** Representative images of the bead-sprouting assay using HUVECs treated with vehicle or ZIETD (10μM) with or without VEGF (50ng/ml) stimulation. **F)** Quantification of the total sprout length shows that blocking Casp-8 activity impairs VEGF-induced sprouting. Approximately 10 beads per condition were quantified, n=4. **G)** BrdU^+^ HUVECs were quantified in control and Casp-8^KD^ ECs with or without VEGF (50ng/ml) or FGF (50 ng/ml) stimulation (24h). Around 100 cells per condition were quantified, n=3. **H)** WST-1 assay showing no response to VEGF-induced proliferation in Casp-8^KD^ ECs (24h), n=9. **I)** Representative bright field images of the scratch migration assay of HUVECs transfected with siCtrl or siCasp-8. Wound closure is reduced in Casp-8^KD^ ECs after 12h of VEGF (50ng/ml) stimulation. **J)** Quantification of the gap closure from (I). 15 fields per condition were quantified, n=5. For C,F-H,J data represent mean ± SEM (* *P* <0.05, ** *P* <0.01, *** *P* <0.001; two-way ANOVA with Bonferroni’s multiple comparisons test). Scale bars: B,E) 100μm; I) 200μm.

**Suppl Fig. 5. Casp-8^ECko^ mice show an active VE-cadherin staining at the junctions of the sprouting front.**

**A)** Representative images of the front of the retina stained with IsoB4 and VE-cadherin. Images of the VE-cadherin single channel were transformed to grey colors with ImageJ for better visualization. **B)** Quantification of the percentage of VE-cadherin patches in the sprouting front showing only a mild reduction in Casp-8^ECko^ mice in the number of highly inhibited VE-cadherin patches compared to Casp-8^WT^. Each box shows the median percentage of patches of that type (line), and upper and lower quartiles (box). The whiskers extend to the most extreme data within 1.5 times the interquartile range of the box. (** *P* <0.01; Dirichlet regression model with two-tailed Mann Whitney test for each state; n=9 WT, 9 ECko). **C)** Average of the differential distribution of the percentage of VE-Cadherin patches of Casp-8^WT^ and Casp-8^ECko^ retinas. Data from 3 independent litters. Scale bars: A) 20μm.

**Suppl Fig. 6. VEGF-induced phosphorylation of Akt, ERK, FAK and p38 is not affected by loss of Casp-8 in ECs.**

**A)** Western blot showing that the Akt, ERK, FAK and p38 signaling pathways are normally activated upon VEGF stimulation in Casp-8^KD^ ECs. **B-E)** Quantification of western blots shown in (A), n=3-5. **F)** Western blot of HUVECs treated with ZIETD (10μM, 16h) and stimulated with VEGF (50ng/ml) for the indicated times, showing that blocking Casp-8 activity also induces increased basal p-p38. Insets of p-p38 at basal conditions (of the same blots) are shown in the upper panels. **G)** Quantification of p-p38 after VEGF stimulation, n=3. **H)** Bright field representative images of the tube formation assay quantified in Fig. 6H. ShCtrl or shCasp-8 infected HUVECs were treated with or without p38 inhibitor (SB) and with VEGF (50ng/ml) for 4h. For B-D, F,H data represent mean ± SEM (ns: not significant; repeated measures two-way ANOVA). Scale bars: I) 100μm.

**Suppl Fig. 7. Knockdown of RIPK3 alone has no impact on angiogenesis.**

**A)** Western blot showing that the sole siRNA-mediated knockdown of RIPK3 in HUVECs does not affect p38 activation. HUVECs were stimulated with VEGF for the indicated time points. **B)** Quantification of western blots shown in (A), n=3. **C)** Bright field representative images of the tube formation assay quantified in Fig. 7D. ShCtrl or shCasp-8 infected HUVECs were transfected with control or Ripk3 siRNA and treated with VEGF (50ng/ml) for 4h. **D,E)** Representative images of whole mount P6 retinas of RIPK3^WT^, heterozygous (RIPK3^Het^) and RIPK3^ko^ mice stained with IsoB4 (ECs) (D) and higher magnifications of the retina front (E). Black dashed lines highlight the total retina area. **F,G)** Quantification of vessel area (F; n=5 WT, 10 Het, 5 ko) and number of branches (G; n=5 WT, 10 Het, 5 ko). Data represent mean ± SEM (ns: not significant; for B repeated measures two-way ANOVA; for F,G one-way ANOVA with Bonferroni’s multiple comparisons test, data from 3 independent litters). Scale bars: C-E) 100μm.

**Suppl. Fig. 8: Vascular defects in Casp-8^ECko^ mice are recovered at P15 and P42.**

**A)** Scheme of the applied extended tamoxifen treatment protocol. **B,C)** Neither the vessel area (B), nor the number of branches (C) are affected at P15 in Casp-8^ECko^ mice (n=6WT, 4ECko). **D)** Retro-orbital injection of 70kDa fluorescently labeled Dextran does not reveal qualitative differences in vessel permeability in neither the superficial, intermediate or deep vascular layers in Casp-8^ECko^ mice compared to wildtype littermates. Representative pictures show Dextran localized inside, but not outside of IsoB4 labeled vessels. **E-G)** Consistently, no defects regarding vessel area (E), number of branches (F) or vessel permeability (G) were observed in Casp-8^ECko^ mice at P42 (n=5WT, 4ECko). For B,C,E,F data represent mean ± SEM (* *P* <0.05, ** *P* <0.01, *** *P* <0.001; two-way ANOVA with Bonferroni’s multiple comparisons test) Scale bars: D,G) 100μm.

