## Supplementary material for "Caspase-8 Modulates Angiogenesis By Regulating A Cell Death Independent Pathway In Endothelial Cells": Suppl. Figures

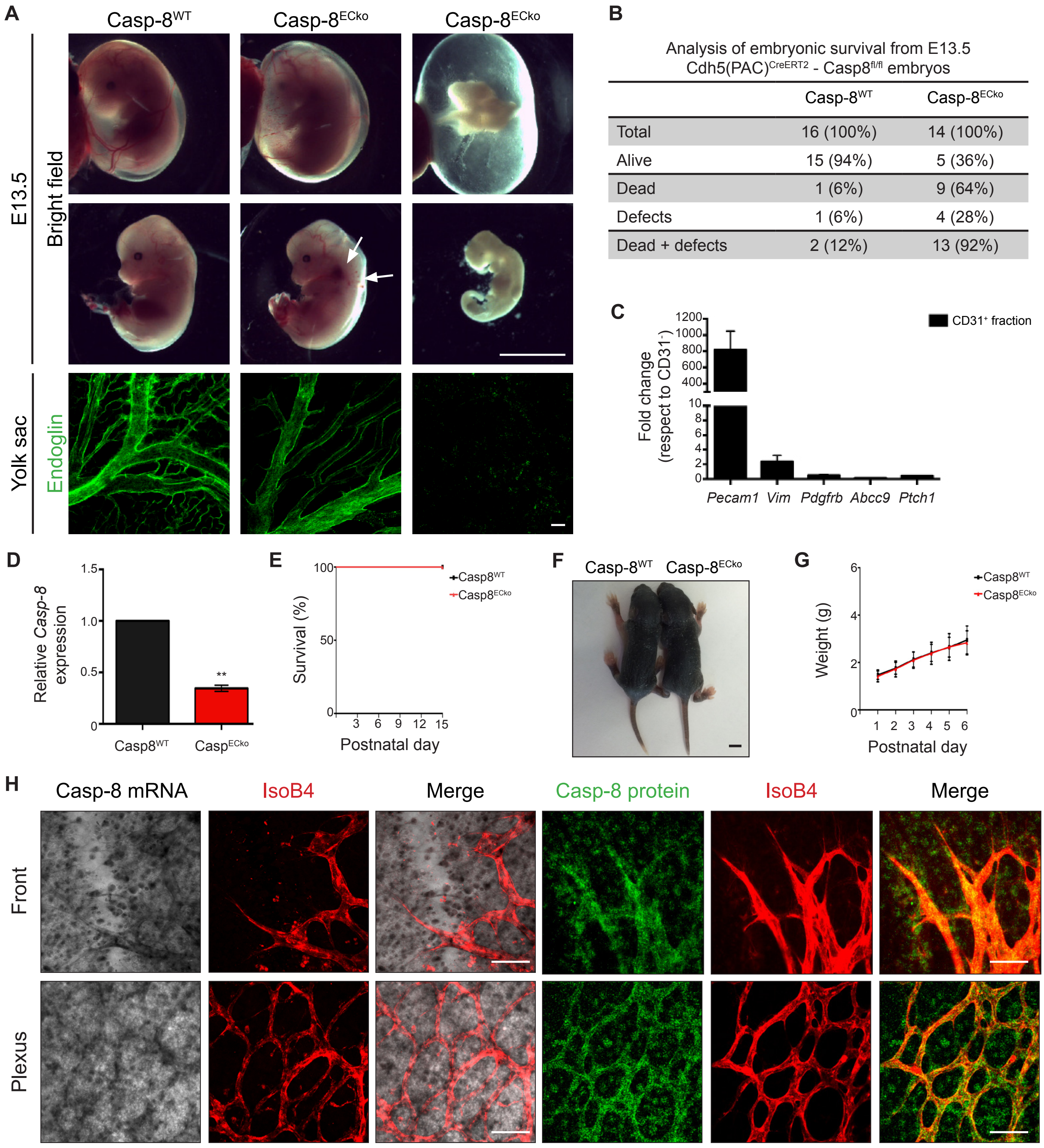


**Suppl Fig 1. Knockout of Casp-8 in ECs is lethal during embryonic, but not postnatal development.**

**A)** Casp-8$^{WT}$ and Casp-8$^{ECko}$ mouse embryos at E13.5. Upper and middle panels: embryo overview showing developmental arrest or severe haemorrhages (white arrows) in Casp-8$^{ECko}$ embryos. Lower panels: immunostaining for the EC marker Endoglin, showing impaired yolk sac vasculature in Casp-8$^{ECko}$ embryos (E13.5). **B)** Table of embryonic lethality of Casp-8$^{ECko}$ embryos at E13.5. **C)** qRT-PCR of isolated lung ECs showing purity of the CD31$^{+}$ fraction. Values were normalized to the CD31$^{-}$ fraction. **D)** qRT-PCR of isolated lung ECs showing knockout efficiency in Casp-8$^{ECko}$ pups (results from 3 independent litters; ** $P$ <0.01; one-sample *t*-test). **E)** No differences between genotypes were observed on mouse weight (n=25 WT, 22 ECko). **F)** Representative images from Casp-8$^{WT}$ and Casp-8$^{ECko}$ mice. **G)** No differences between genotypes were observed on survival (n=25 WT, 22 ECko). **H)** *In situ* hybridization (mRNA) or immunofluorescence (protein) showing Casp-8 expression at the front and the back (plexus) of P6 mouse retinas. Data represent mean ± SEM. Scale bars: A) 1cm (upper panel) and 100μm (lower panel); F) 5mm; H) 100μm.

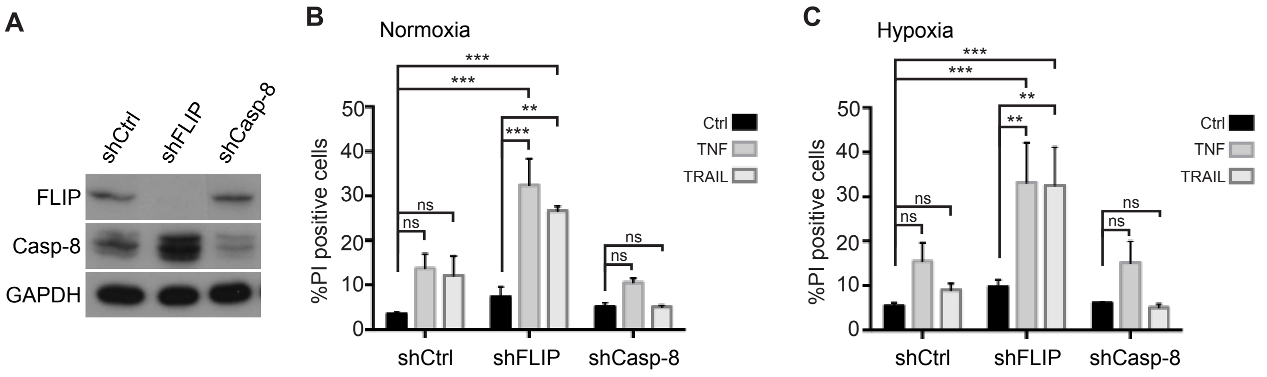


**Suppl Fig. 2. Knockdown of Casp-8 in ECs does not increase cell death *in vitro*.**

**A)** Western blot showing protein knockdown efficiency after infecting HUVECs with a lentivirus carrying shRNAs for c-FLIP or Casp-8. **B,C)** Quantification of apoptotic/necrotic HUVECs ($PI^+$) in normoxia (B) or hypoxia (C) showing that knockdown of c-FLIP (shFLIP), after the indicated treatments, increases cell death while knockdown of Casp-8 (shCasp-8) does not, n=3. Data represent mean ± SEM (** $P$ <0.01, *** $P$ <0.001, ns: not significant; two-way ANOVA with Bonferroni's multiple comparisons test).

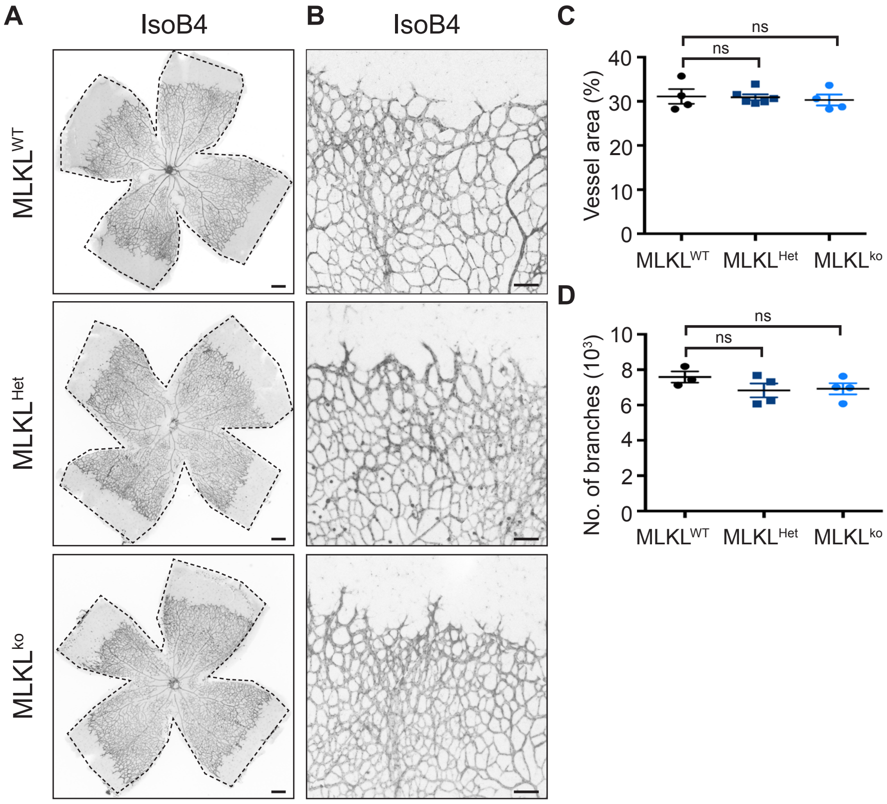


**Suppl Fig. 3. MLKL knockout pups do not present impaired postnatal angiogenesis.** **A,B)** Representative images of whole mount P6 retinas of MLKL$^{WT}$, MLKL heterozygous (MLKL$^{Het}$) and MLKL$^{ko}$ mice stained with IsoB4 (ECs) (A) and higher magnifications of the retina front (B). Black dashed lines highlight the total retina area. **C,D)** Quantification of vessel area (C; n=4 WT, 6 Het, 4 ko) and number of branches (D; n=3 WT, 4 Het, 4 ko). For C,D data represent mean ± SEM from 2 independent litters (ns: not significant; one-way ANOVA with Bonferroni's multiple comparisons test). Scale bars: A) 100µm; B) 50µm.

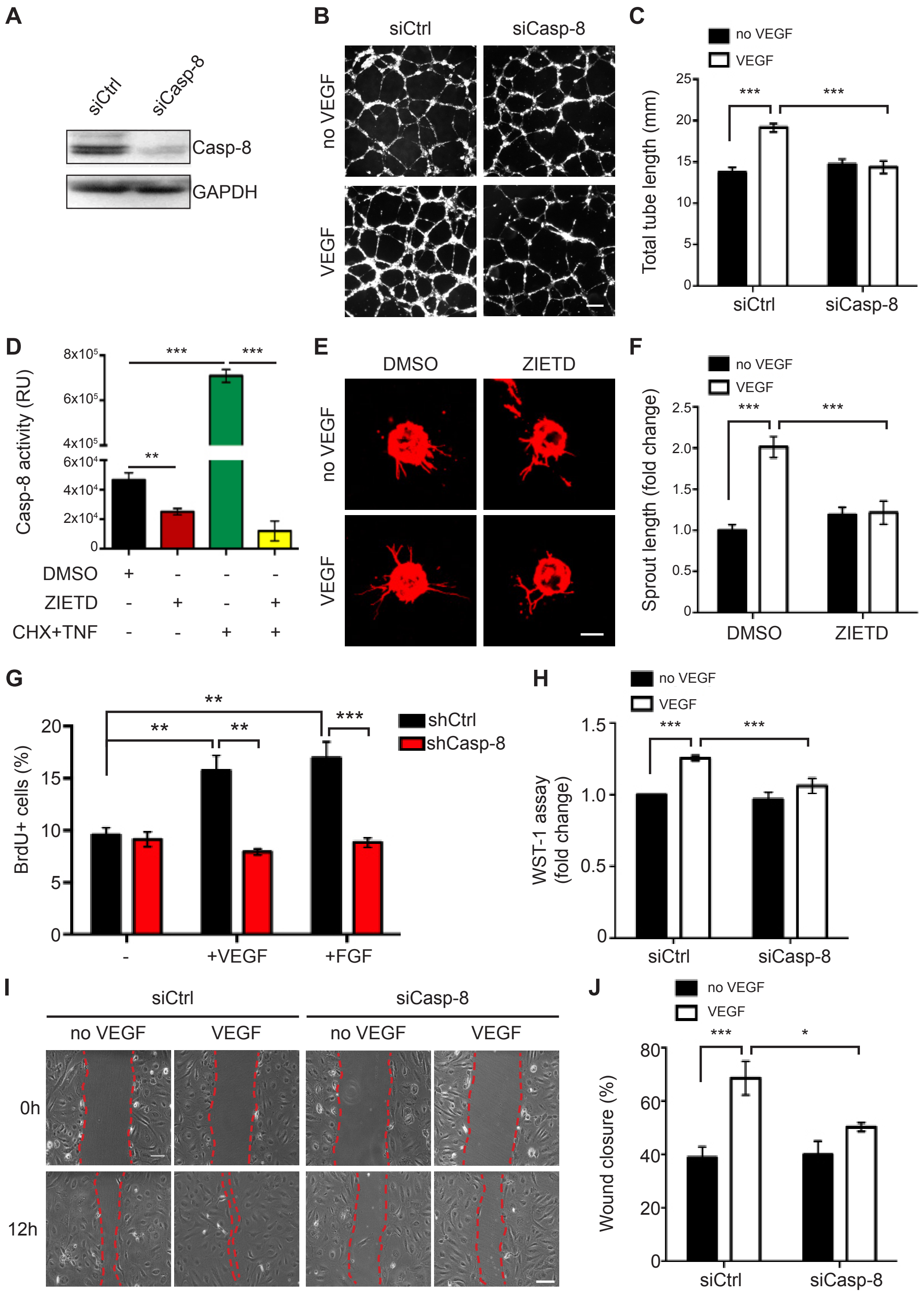


**Suppl Fig. 4. Knockdown of Casp-8 impairs VEGF-induced tube formation, EC proliferation and migration *in vitro.***
**A)** Western blot showing the knockdown efficiency of the Casp-8 siRNA. **B)** Bright field representative images of the tube formation assay. HUVECs were transfected with control siRNA (siCtrl) or Casp-8 siRNA (siCasp-8) and treated with VEGF (50ng/ml) for 4h. **C)** Quantification of the total tube length as in (B) showing that Casp-8 knockdown inhibits VEGF-induced tube formation. 5 fields per condition were quantified, n=3. **D)** Casp-8 activity was measured with a Casp-8 Glo kit. HUVECs' basal Casp-8 activity can be blocked with the Casp-8 inhibitor ZIETD (10µM) or induced by cycloheximide (CHX, 1µg/ml) plus TNF (100ng/ml) for 4h, n=3. Data represent mean ± SEM (** $P$ <0.01, *** $P$ <0.001; two-tailed unpaired Student *t*-test). **E)** Representative images of the bead-sprouting assay using HUVECs treated with vehicle or ZIETD (10µM) with or without VEGF (50ng/ml) stimulation. **F)** Quantification of the total sprout length shows that blocking Casp-8 activity impairs VEGF-induced sprouting. Approximately 10 beads per condition were quantified, n=4. **G)** BrdU$^+$ HUVECs were quantified in control and Casp-8$^{KD}$ ECs with or without VEGF (50ng/ml) or FGF (50 ng/ml) stimulation (24h). Around 100 cells per condition were quantified, n=3. **H)** WST-1 assay showing no response to VEGF-induced proliferation in Casp-8$^{KD}$ ECs (24h), n=9. **I)** Representative bright field images of the scratch migration assay of HUVECs transfected with siCtrl or siCasp-8. Wound closure is reduced in Casp-8$^{KD}$ ECs after 12h of VEGF (50ng/ml) stimulation. **J)** Quantification of the gap closure from (I). 15 fields per condition were quantified, n=5. For C,F-H,J data represent mean ± SEM (* $P$ <0.05, ** $P$ <0.01, *** $P$ <0.001; two-way ANOVA with Bonferroni's multiple comparisons test). Scale bars: B,E) 100µm; I) 200µm.

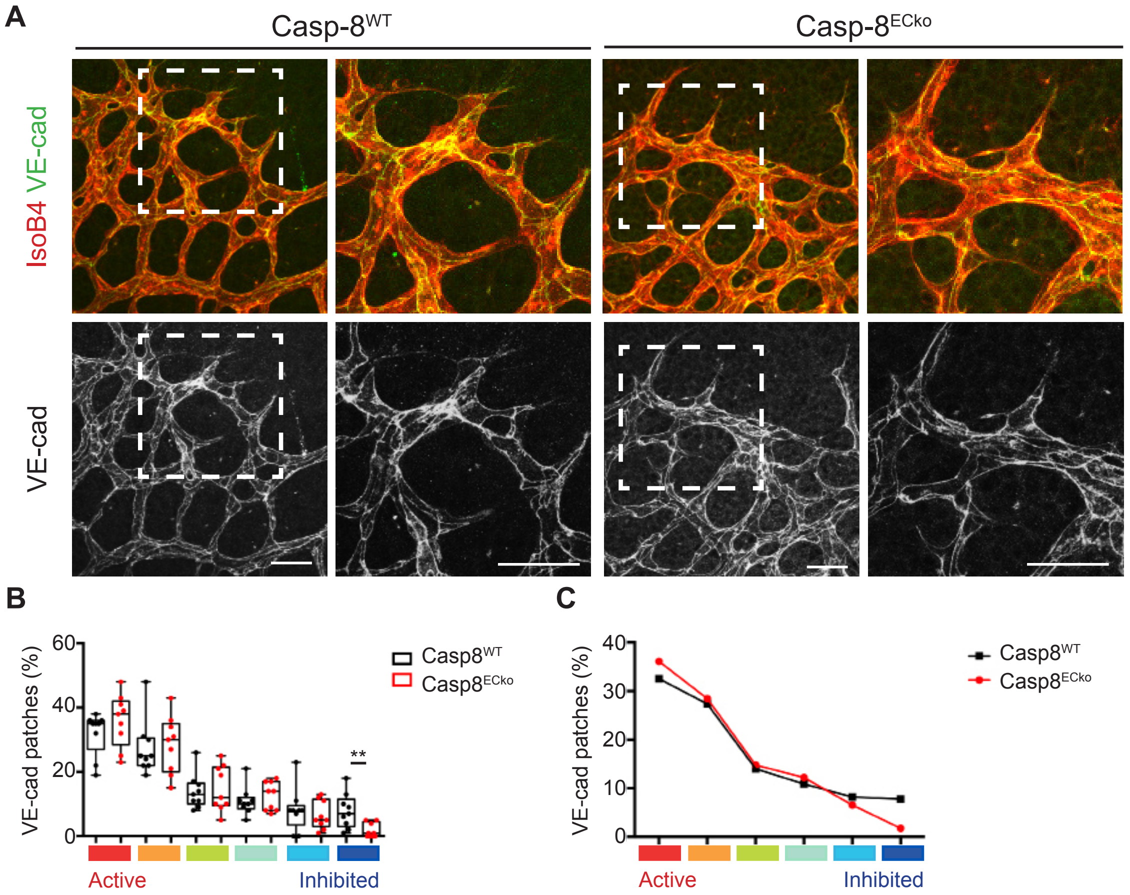


**Suppl Fig. 5. Casp-8$^{ECko}$ mice show an active VE-cadherin staining at the junctions of the sprouting front.**
**A)** Representative images of the front of the retina stained with IsoB4 and VE-cadherin. Images of the VE-cadherin single channel were transformed to grey colors with ImageJ for better visualization. **B)** Quantification of the percentage of VE-cadherin patches in the sprouting front showing only a mild reduction in Casp-8$^{ECko}$ mice in the number of highly inhibited VE-cadherin patches compared to Casp-8$^{WT}$. Each box shows the median percentage of patches of that type (line), and upper and lower quartiles (box). The whiskers extend to the most extreme data within 1.5 times the interquartile range of the box. ( ** *P* <0.01; Dirichlet regression model with two-tailed Mann Whitney test for each state; n=9 WT, 9 ECko). **C)** Average of the differential distribution of the percentage of VE-Cadherin patches of Casp-8$^{WT}$ and Casp-8$^{ECko}$ retinas. Data from 3 independent litters. Scale bars: A) 20µm.

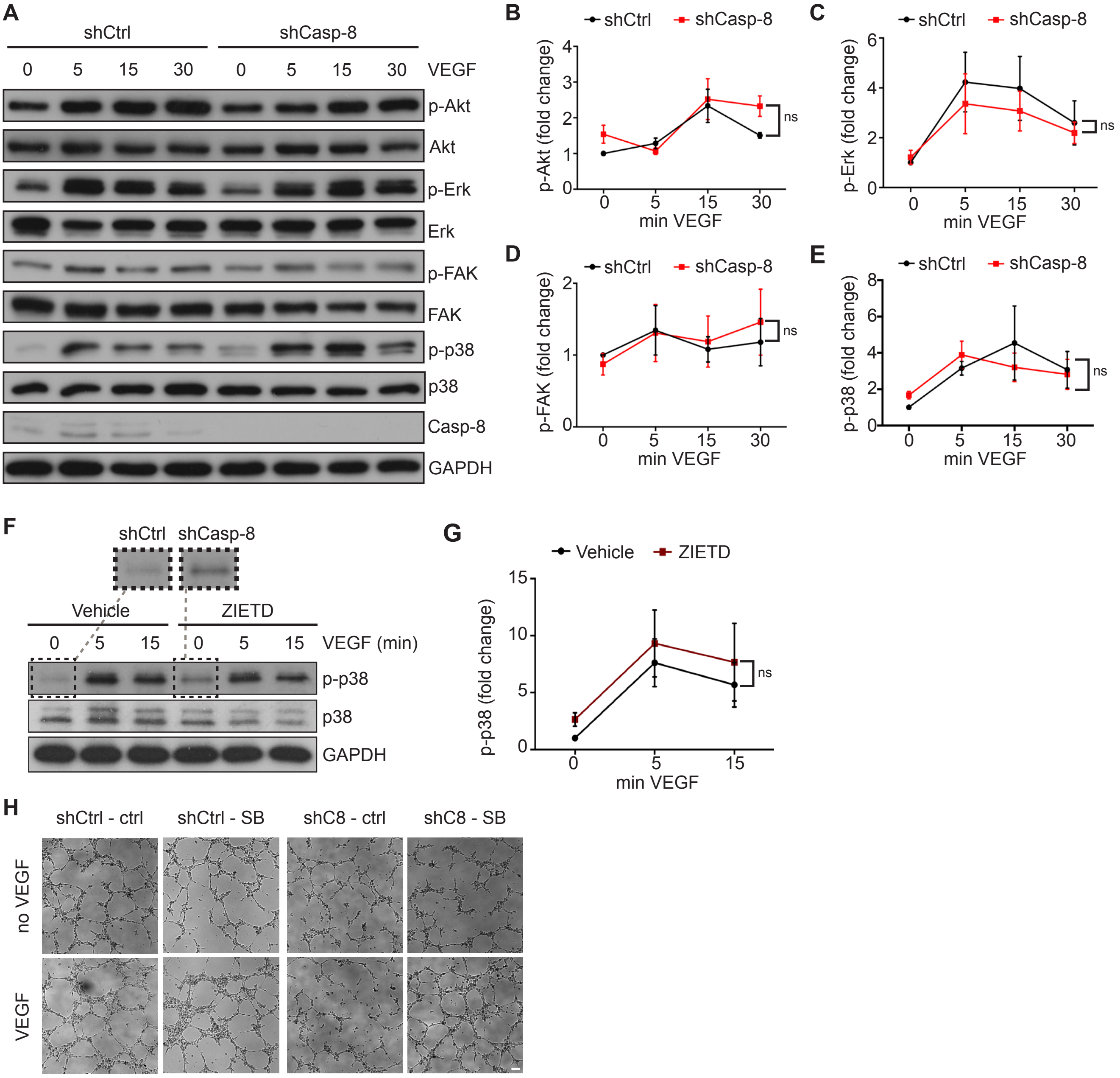


**Suppl Fig. 6. VEGF-induced phosphorylation of Akt, ERK, FAK and p38 is not affected by loss of Casp-8 in ECs.**
**A)** Western blot showing that the Akt, ERK, FAK and p38 signaling pathways are normally activated upon VEGF stimulation in Casp-8$^{KD}$ ECs. **B-E)** Quantification of western blots shown in (A), n=3-5. **F)** Western blot of HUVECs treated with ZIETD (10µM, 16h) and stimulated with VEGF (50ng/ml) for the indicated times, showing that blocking Casp-8 activity also induces increased basal p-p38. Insets of p-p38 at basal conditions (of the same blots) are shown in the upper panels. **G)** Quantification of p-p38 after VEGF stimulation, n=3. **H)** Bright field representative images of the tube formation assay quantified in Fig. 6H. ShCtrl or shCasp-8 infected HUVECs were treated with or without p38 inhibitor (SB) and with VEGF (50ng/ml) for 4h. For B-D, F,H data represent mean ± SEM (ns: not significant; repeated measures two-way ANOVA). Scale bars: I) 100µm.

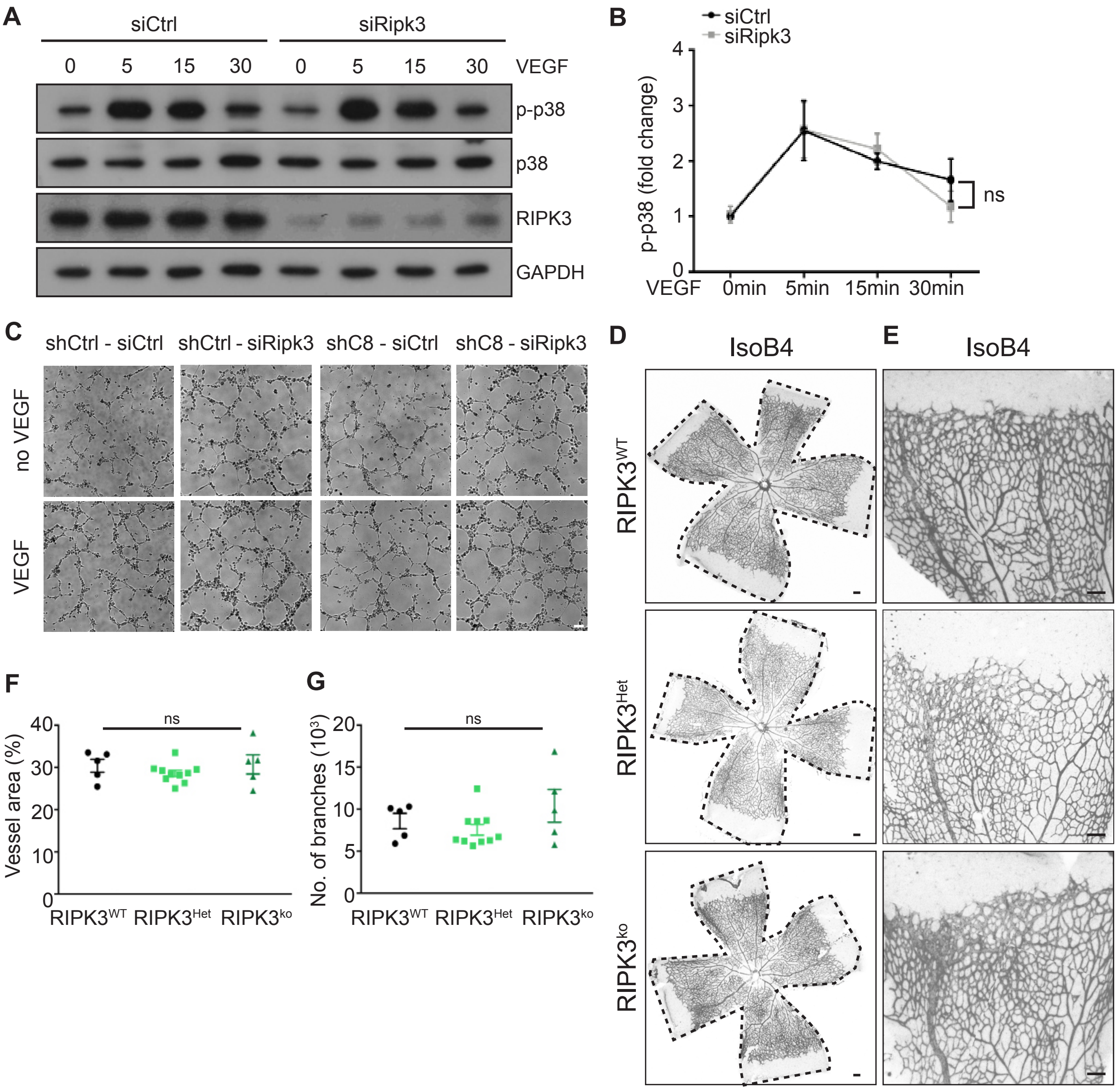


**Suppl Fig. 7. Knockdown of RIPK3 alone has no impact on angiogenesis.**

**A)** Western blot showing that the sole siRNA-mediated knockdown of RIPK3 in HUVECs does not affect p38 activation. HUVECs were stimulated with VEGF for the indicated time points. **B)** Quantification of western blots shown in (A), n=3. **C)** Bright field representative images of the tube formation assay quantified in Fig. 7D. ShCtrl or shCasp-8 infected HUVECs were transfected with control or Ripk3 siRNA and treated with VEGF (50ng/ml) for 4h. **D,E)** Representative images of whole mount P6 retinas of RIPK3$^{WT}$, heterozygous (RIPK3$^{Het}$) and RIPK3$^{ko}$ mice stained with IsoB4 (ECs) (D) and higher magnifications of the retina front (E). Black dashed lines highlight the total retina area. **F,G)** Quantification of vessel area (F; n=5 WT, 10 Het, 5 ko) and number of branches (G; n=5 WT, 10 Het, 5 ko). Data represent mean ± SEM (ns: not significant; for B repeated measures two-way ANOVA; for F,G one-way ANOVA with Bonferroni's multiple comparisons test, data from 3 independent litters). Scale bars: C-E) 100μm.

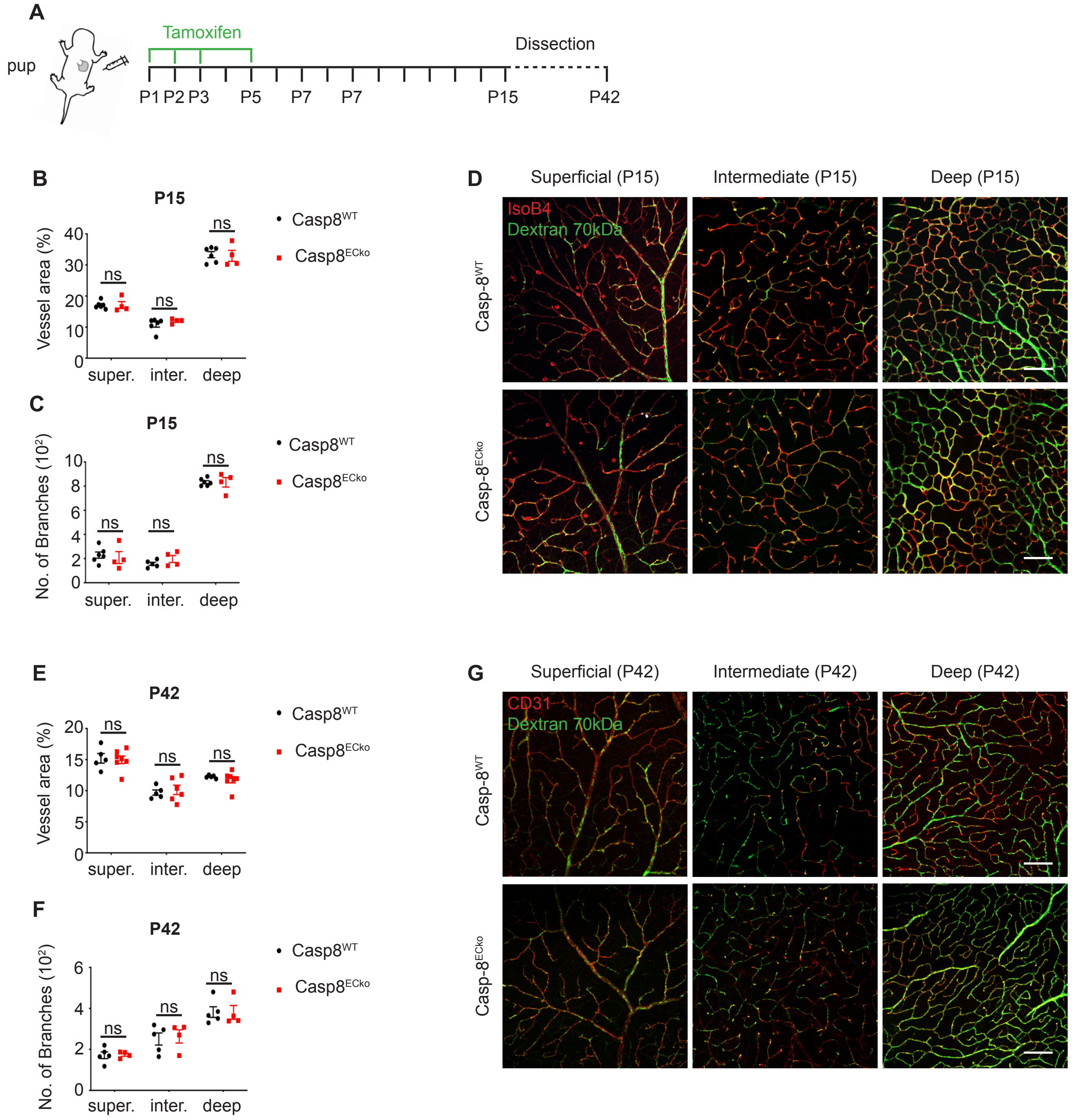


**Suppl. Fig. 8: Vascular defects in Casp-8$^{ECko}$ mice are recovered at P15 and P42.**

**A)** Scheme of the applied extended tamoxifen treatment protocol. **B,C)** Neither the vessel area (B), nor the number of branches (C) are affected at P15 in Casp-8$^{ECko}$ mice (n=6WT, 4ECko). **D)** Retro-orbital injection of 70kDa fluorescently labeled Dextran does not reveal qualitative differences in vessel permeability in neither the superficial, intermediate or deep vascular layers in Casp-8$^{ECko}$ mice compared to wildtype littermates. Representative pictures show Dextran localized inside, but not outside of IsoB4 labeled vessels. **E-G)** Consistently, no defects regarding vessel area (E), number of branches (F) or vessel permeability (G) were observed in Casp-8$^{ECko}$ mice at P42 (n=5WT, 4ECko). For B,C,E,F data represent mean ± SEM (* *P* <0.05, ** *P* <0.01, *** *P* <0.001; two-way ANOVA with Bonferroni's multiple comparisons test) Scale bars: D,G) 100µm.
